# A synthetic morphogenic membrane system that responds with self-organized shape changes to local light cues

**DOI:** 10.1101/481887

**Authors:** Konstantin Gavriljuk, Bruno Scocozza, Farid Ghasemalizadeh, Akhilesh P. Nandan, Manuel Campos Medina, Hans Seidel, Malte Schmick, Aneta Koseska, Philippe I. H. Bastiaens

**Affiliations:** Department of Systemic Cell Biology, Max Planck Institute for Molecular Physiology, 44227 Dortmund, Germany; Faculty of Chemistry and Chemical Biology, TU Dortmund, 44227 Dortmund, Germany

**Keywords:** Synthetic proto-cell, cell morphogenesis, reconstituted microtubule and signaling system, dimensionality reduction, signaling gradient, membrane-cytoskeletal interactions, cellular information processing, morphological plasticity, self-organization

## Abstract

Reconstitution of artificial cells capable of transducing extracellular signals into cytoskeletal changes is a challenge in synthetic biology that will reveal fundamental principles of non-equilibrium phenomena of cellular morphogenesis and information processing. Here, we generated a ‘life-like’ Synthetic Morphogenic Membrane System (SynMMS) by encapsulating a dynamic microtubule (MT) aster and a light-inducible signaling system driven by GTP/ATP chemical potential into cell-sized vesicles. The biomimetic design of the light-induced signaling system embodies the operational principle of morphogen induced Rho-GTPase signal transduction in cells. Activation of synthetic signaling promotes membrane-deforming growth of MT-filaments by dynamically elevating the membrane-proximal concentration of tubulin. The resulting membrane deformations enable the recursive coupling of the MT-aster with the signaling system, creating global self-organized morphologies that reorganize towards external light cues in dependence on prior sensory experience that is stored in the dynamically maintained morphology. SynMMS thereby signifies a step towards bio-inspired engineering of self-organized cellular morphogenesis.

## INTRODUCTION

Eukaryotic cells have evolved a broad range of solutions for gathering, storing and processing extracellular information to adapt or maintain their morphological identity in a changing environment. Cells acquire their shape, which is tightly linked to their biological function, by dynamic cytoskeletal systems that deform the plasma membrane. Cell shape changes and motility are driven by the microtubule (MT) and actin filament systems, which operate at different length- and time-scales^1^. Actin filaments dictate the rapid, more local morphological dynamics at the cell periphery, whereas MTs are far more long lived, persist over longer distances and are globally organized through MT organizing centers (MTOC) during developmental processes. MTs thereby function as a single cohesive network on the scale of the cell that not only operates as a compass for front-back polarity in cell movement, but also generates and maintains global cell shapes during differentiation^2–6^. The dynamical organization of the MT network must therefore account for both, plasticity in shape formation in undifferentiated cells depending on environmental cues, but also shape stabilization after differentiation. In these morphogenic processes, signal transduction networks are fundamental and allow cells to map outside information into an inner biochemical state that induces reorganization of their shape, e.g., by morphogen-induced Rho-GTPase signaling that affects cytoskeletal dynamics by regulation of microtubule-associated proteins (MAPs).

However, cells do not reorganize their shape solely in response to extracellular signals, but integrate in their response previous sensory experiences in order to commit to distinct shapes during differentiation^7–9^. How memory of previous morphogen patterns is realized and affects signal-induced morphogenesis remains elusive. Based on prior work relating local geometric features of the plasma membrane to signaling activity^10^, we have postulated that a closed loop causality between membrane shape, signalling and cytoskeletal dynamics underlies cellular morphological responses^11^. How such cellular functionalities emerge from the dynamics of interacting sub-systems can be experimentally addressed in synthetic proto-cells where the functional molecular modules are encapsulated under non-equilibrium conditions^12^. Previous reconstitutions, although mainly focused on the interplay between the MT-cytoskeleton and motor proteins, have already revealed much about the fundamental mechanisms of self-organized cell polarity^13–19^.

To capture ubiquitous mechanisms of cellular information processing that translate into cellular morphogenesis, we designed a minimal system in cell-sized liposomes capable of transducing external stimuli into internal signaling gradients that affect MT-aster dynamics^20^. The light-activatable synthetic signaling system mimics the dimensionality reduction^21–23^ operational principle of the canonical Rac1-Pak1-stathmin MT-regulator pathway^24–27^ (Fig. 1) and the directional response to an extracellular morphogen gradient. Our results demonstrate that signaling-induced membrane deformations not only define cell shape, but also constitute a means by which the MT-cytoskeleton and signaling can recursively interact, causing these proto-cells to self-organize into shapes that can be transformed and stabilized upon extracellular cues. A reaction-diffusion theoretical framework of the coupled cytoskeletal and signaling dynamics captured the initial morphological states as well as light-induced morphological transitions. We thereby show that the interdependence of MT dynamics and signaling as mediated by the deformable membrane is a fundamental mechanism by which cells utilize previous sensory experiences in reorganizing their shape in response to extracellular signals.

**Fig. 1.**
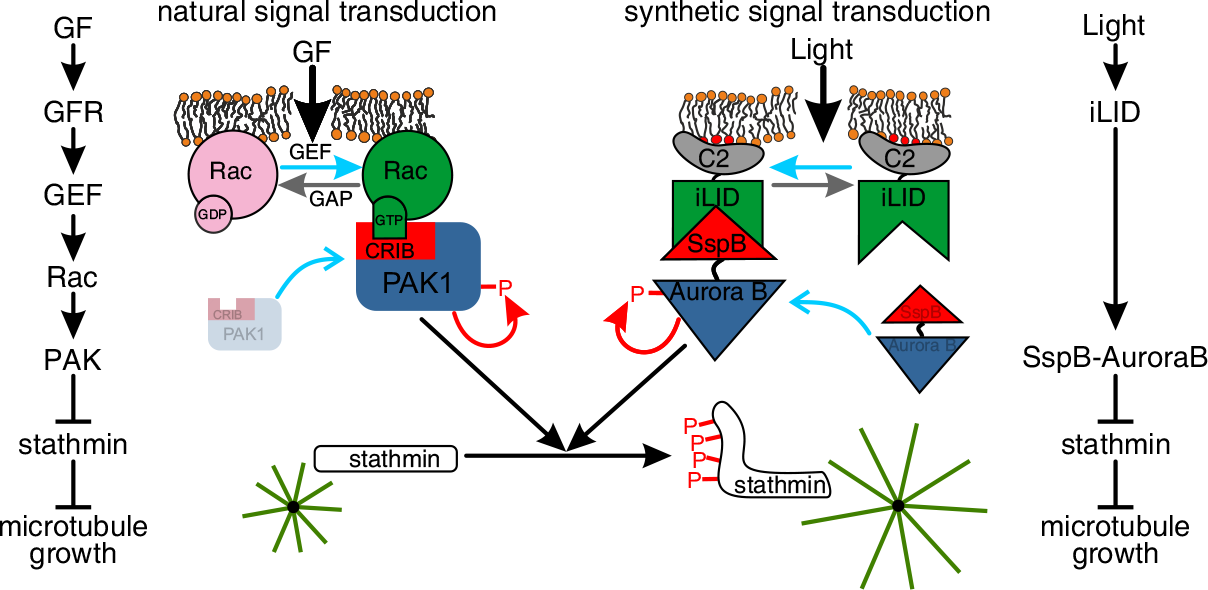
Synthetic network mimics natural Rho-GTPase mediated signal transduction. Natural (left) and synthetic (right) signaling pathways. Center: corresponding activation (vertical black arrows) of natural or synthetic signal transduction by translocation of PAK to Rac·GTP (via CRIB domain) or SspB-AuroraB to C2-iLID on the membrane (curved blue arrows). Upon activation of kinase by autophosphorylation (red arrows) MT growth increases (green asters) by phosphorylation of stathmin (horizontal black arrow). GF: growth factor, GFR: growth factor receptor, GEF: guanine nucleotide exchange factor, GAP: GTPase activating protein, Rac: small GTPase, PAK, AuroraB: serine threonine protein kinases, stathmin: MT regulator, iLID/SspB: improved light-inducible dimer and its binding partner, C2: phosphatidylserine binding domain.

## RESULTS

### The MT-asters membrane system

To first investigate which morphologies emerge from the collective behavior of centrally organized dynamic MTs that interact via a deformable membrane, we encapsulated purified centrosomes together with tubulin and GTP in GUVs using cDICE^28^, and controlled the deformability of the membrane by the outside osmolarity (Methods). This generated dynamically instable^29^ MT-asters with different sizes inside the GUVs, which could be monitored by confocal laser scanning microscopy (CLSM) using trace amounts of fluorescently labeled tubulin (~10 % Alexa568- or Alexa488-tubulin, Methods). Encapsulating low tubulin concentrations (15 – 25 μM) under isosmotic conditions resulted in GUVs with a spherical morphology and centrally positioned asters whose size was smaller than that of the GUV, while higher tubulin concentrations (35 – 40 μM) resulted in cortical MTs with peripheral centrosome positioning (Fig. 2a,b; Supplementary Fig. 1a,b)^18^. Decreasing the membrane tension on the other hand, led to a polar morphology with one or more protrusions, reflected by an increased GUV eccentricity (Fig. 2 a,b). The latter morphologies were maintained by the dynamics of the MT-asters, as apparent from the morphological transition from spherical to polar protrusion upon increase of the outside osmolarity (Supplementary Fig. 1d,e). This shows that external influences on the dynamics of the MT-membrane system can remodel the morphology.

**Fig. 2.**
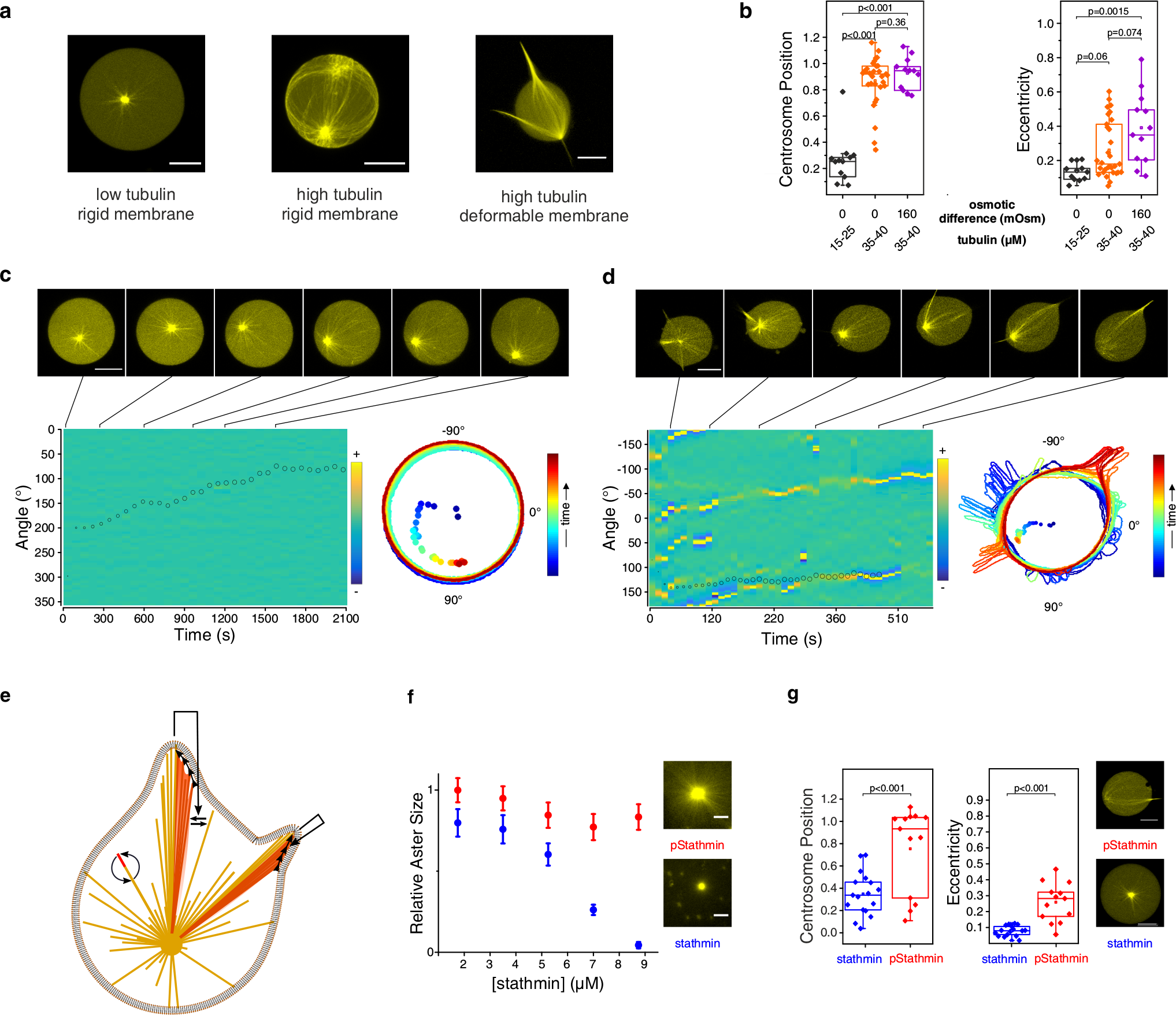
Self-organized polar morphology of encapsulated MT-asters can be controlled by stathmin phosphorylation. (a) GUV morphologies, from left to right: small aster at low tubulin (15 – 25 μM), cortical aster and polar morphologies at high tubulin concentration (35 – 40 μM). Right GUV at lowered membrane tension (hyperosmotic surrounding solution). (b) Morphometric quantification of individual GUVs with high or lowered membrane tension, and low or high tubulin concentration. Centrosome position: 0 – centered, 1 – membrane proximal, eccentricity: 0 – perfect circle, >0 – deviations from circle. (c) Top: time-lapse of temperature-induced aster growth in a GUV with high membrane tension. Bottom left: kymograph of local membrane curvature (right: color code) overlaid with centrosome position (small circle: central, large circle: peripheral). Bottom right: GUV contours during time-lapse, color coded by time progression (dots: corresponding centrosome positions). See also Supplementary Fig. 1g. (d) Time-lapse of temperature-induced aster growth in GUVs with low membrane tension. Representation as in (c). See also Supplementary Fig. 1g. (e) Scheme of self-induced capture (SIC) protrusion formation. MT-induced membrane deformations promote further capture of neighboring MTs (straight arrows), which bundle by sliding into the protrusion (arrow heads) thereby depleting free MTs. Curved arrows represent MT dynamic instability. (F) MT-aster size in solution as a function of stathmin (blue points) and pStathmin (red points) concentration at 35 μM tubulin. Error bars: Standard Error of the regression (Methods). Right: examples of overlaid asters (> 30 asters per condition) at 7 μM pStathmin (top) or stathmin (bottom). (g) Morphometric quantification as in (b) of GUVs containing 5 μM stathmin or pStathmin (40 μM tubulin). Right: representative images. All p-values from two-sample Kolmogorov-Smirnov test. Scale bars 10 μm.

To study whether and which morphological transitions can occur by external regulation of the aster size, the net growth of small encapsulated asters was induced by raising the temperature from 20 °C to 34 °C (Supplementary Fig. 1c). This is analogous to a uniform morphogen signal that globally affects MT-aster size. In GUVs with high membrane tension, temperature-induced MT growth led to centrosome decentering, resulting in the formation of semi-asters ^18^ (Fig. 2c; Supplementary Fig. 1f; Supplementary Video 1). In contrast, in GUVs with a deformable membrane, the system evolved from a morphology with isotropically distributed membrane deformations to a stable polar protrusion morphology. The centrosome decentered as transient protrusions converged to a single one at the opposite pole (Fig. 2d; Supplementary Fig. 1g, Supplementary Video 2). This indicated that growing MT-bundles in protrusions provided an initially random directional push to the centrosome which polarized the GUV by a recursive process of protrusion coalescence and centrosome decentering. The convergence of bending MTs into the protrusions and their coalescence (Fig. 2a,d, Supplementary Fig. 1g) indicates that they were created by self-induced capture (SIC), where MT-induced small membrane deformations served as capture-sites for neighboring microtubules (Fig. 2e). Thus, the indirect coupling between the astral MTs via the deformable membrane gave rise to collective MT behavior resulting in a symmetry broken morphology with protrusions at the poles.

To establish a biochemical regulation of encapsulated asters, we quantified the effects of the MT-regulator stathmin on MT-dynamics, first by single filament TIRF assays (Methods). Increase in stathmin concentration linearly decreased MT-growth speed and abruptly increased catastrophe frequency, whereas phosphorylated stathmin (pStathmin) only weakly affected MT-dynamics (Supplementary Fig. 2a,b). Interaction of fluorescently tagged stathmin with MT plus ends was not observed, concluding that stathmin affects MT-dynamics purely through sequestering free tubulin^20,27,30^. Encapsulation of stathmin (5μM) in GUVs reduced MT-aster size (Fig. 2f,g, Supplementary Fig. 2c), thereby leading to a spherical morphology, whereas encapsulation of 5 μM pStathmin resulted in the polar morphology observed with high tubulin alone (Fig. 2a,g). These results show that MT-induced GUV morphology can be biochemically controlled by a signaling system that phosphorylates stathmin. We therefore reconstituted a signal actuation system based on light-induced kinase translocation to the GUV membrane (Fig. 1).

### The light-induced signaling system

To capture the dimensionality-reduction^21^ principle of Rho GTPase signaling, we encapsulated the iLID/SspB optical dimerizer system^31^ to translocate the stathmin phosphorylating AuroraB kinase^32^ to the membrane. AuroraB was fused to SspB (SspB-AuroraB) and iLID associated with the membrane via a C2 phosphatidylserine-binding domain (C2-iLID) (Fig. 1, right). The amount of SspB-AuroraB translocation could be controlled by blue light intensity (Fig. 3a), where residual binding in the dark determined basal SspB-AuroraB activity and saturatable binding to C2-iLID limited the maximal achievable activity (Supplementary Fig. 3a-c). Importantly, localized blue light irradiation resulted in a translocation gradient along the membrane, showing that the system can mimic a spatially graded response as if originating from an extracellular morphogen gradient (Fig. 3b; Supplementary Fig. 3d,e).

**Fig. 3.**
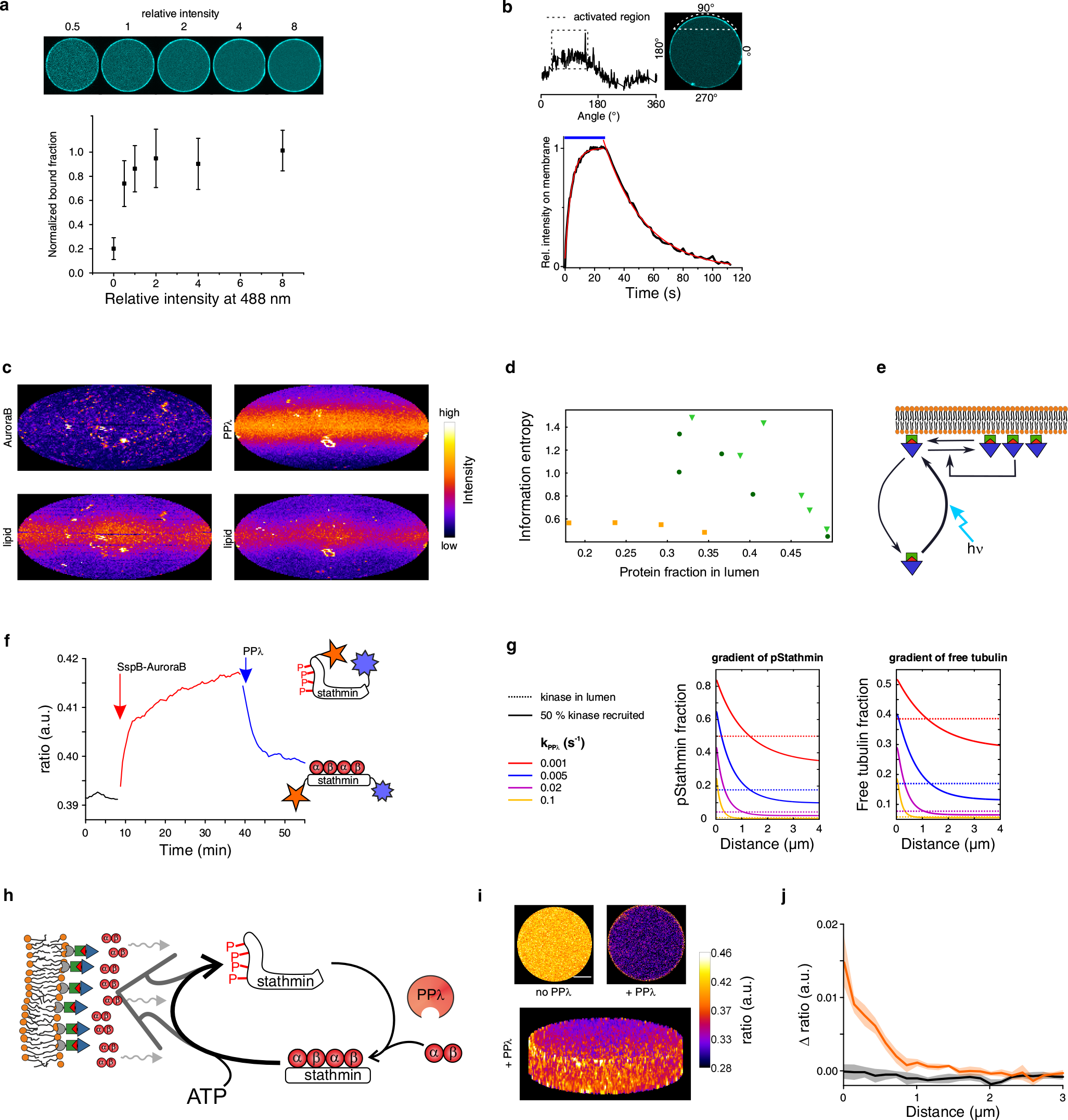
A synthetic light-inducible signaling system generates a membrane-proximal tubulin gradient. (a) Bottom: light-dose response of SspB-AuroraB fraction recruited to the membrane (mean ± S.D. of 5 GUVs, 7 μM C2-iLID, 12 μM Alexa488-SspB-AuroraB). Top: corresponding example images. Laser power normalized to 65 mW/cm^2^. (b) Top: SspB-AuroraB translocation gradient resulting from continuous blue light irradiation of a localized region (white dotted lines). Bottom: kinetics of light-induced membrane recruitment and release in absence of photo-activating light of Alexa647-SspB-AuroraB. Apparent rate constants from exponentially fitting 4 GUVs: k_recruit_ = 13.7 ± 0.3 min^−1^, k_release_ = 2.15 ± 0.06 min^−1^. See also Supplementary Fig. 3d,e. (c) Representative surface intensity maps of translocated SspB-AuroraB (top left) or SspB-PPλ (top right) in GUVs. Bottom: corresponding fluorescent LissamineRhodamineB-DOPE lipid distribution. See also Supplementary Fig. 3f. (d) Respective recurrence quantification analysis of protein spatial pattern regularity (Methods). Information entropy as function of protein depletion from lumen in individual GUVs is shown. Dark green: SspB-AuroraB with ATP, light green: without ATP, orange: SspB-PPλ. See also Supplementary Fig. 3g. (e) Scheme of SspB-AuroraB (blue triangles) organization mechanism based on light-induced substrate-depletion (curved arrows) and cooperative (feedback arrow) clustering (horizontal arrows). (f) Ratiometric measurement of sequential COPY° phosphorylation and dephosphorylation in bulk (10 μM COPY°, 20 μM tubulin, 2 μM SspB-AuroraB, 500 nM PPλ). Enzyme addition indicated by arrows. Right: scheme of the FRET-sensor COPY° (stathmin conjugated to Atto532 (orange) and Atto655 (violet)), in which phosphorylation induced release of αβ-tubulin heterodimers (red circles) causes a conformational change leading to increased FRET efficiency. (g) 1D reaction-diffusion simulations of stathmin phosphorylation (left) and free tubulin (right) profiles upon rebalancing phosphorylation/dephosphorylation cycles by kinase translocation. Profiles are plotted from the membrane (0 μm) before (kinase only in lumen with k_kin_ = 0.004 s^−1^, dashed lines) and after kinase recruitment (50 % kinase on membrane with k_kin_ = 1 s^−1^, solid lines) for varying k_PPλ_ as indicated in the legend (left)(see Methods). (h) Stathmin phosphorylation cycle maintains an enhanced tubulin concentration near the membrane by AuroraB kinase (blue triangles) mediated release of tubulin (red circles) from phosphorylated stathmin (dark gray arrow towards membrane), which is countered by tubulin diffusion (wiggly arrows). Diffusing phosphorylated stathmin is dephosphorylated by luminal PPλ, to rebind tubulin hetero-dimers, closing the ATP driven tubulin deposition cycle. (i) COPY° phosphorylation gradient inside GUVs. Top: maximum intensity projections of 8-slice ratiometric z-stacks without (left) and with PPλ (right), bottom: 3D-projection of the ratiometric z-stack with PPλ. See also Supplementary Fig. 5. (j) Ratiometric, baseline subtracted profiles of COPY° fluorescence (Δratio) inside GUVs. Without PPλ (black): mean ± S.E.M. of 6 GUVs, with PPλ (orange): mean ± S.E.M. of 4 GUVs. Scale bars 10 μm.

We observed that SspB-AuroraB clusters were formed on the membrane upon its light-induced translocation, irrespective of ATP, and therefore kinase activity. This clustering was a specific property of AuroraB and not of the iLID/SspB proteins, since it did not occur for a SspB-λ-phosphatase fusion construct (SspB-PPλ) (Fig. 3c; Supplementary Fig. 3f). Quantification of the spatial SspB-AuroraB pattern recurrences on the membrane by means of information entropy^33^ (Methods) revealed that the spatial order increased for higher SspB-AuroraB depletion from the lumen, which was not observed for the spatial SspB-PPλ distribution (Fig. 3d; Supplementary Fig. 3g). This indicates that light-induced translocation drives self-organized SspB-AuroraB pattern formation based on substrate-depletion, thereby generating a persistent and patterned response (Fig. 3e).

We next introduced stathmin in the system to couple SspB-AuroraB kinase translocation to MT-dynamics regulation. We first measured bulk solution kinetics of stathmin phosphorylation by SspB-AuroraB and dephosphorylation by PPλ (Supplementary Table 1) in the presence of soluble tubulin (20 μM), using an organic dye-containing variant of the stathmin phosphorylation FRET sensor COPY^20^ (Atto532-stathmin-Atto655: COPY° (organic)) (Supplementary Fig. 4a,b). Ratiometric quantification COPY° phosphorylation after sequential addition of SspB-AuroraB and PPλ at equal final concentration (1 μM), demonstrated that phosphorylation cycles can maintain a low steady state phosphorylation level of stathmin in the presence of ATP (Fig. 3f; Supplementary Fig. 4c,d; Supplementary Table 1).

**Fig. 4.**
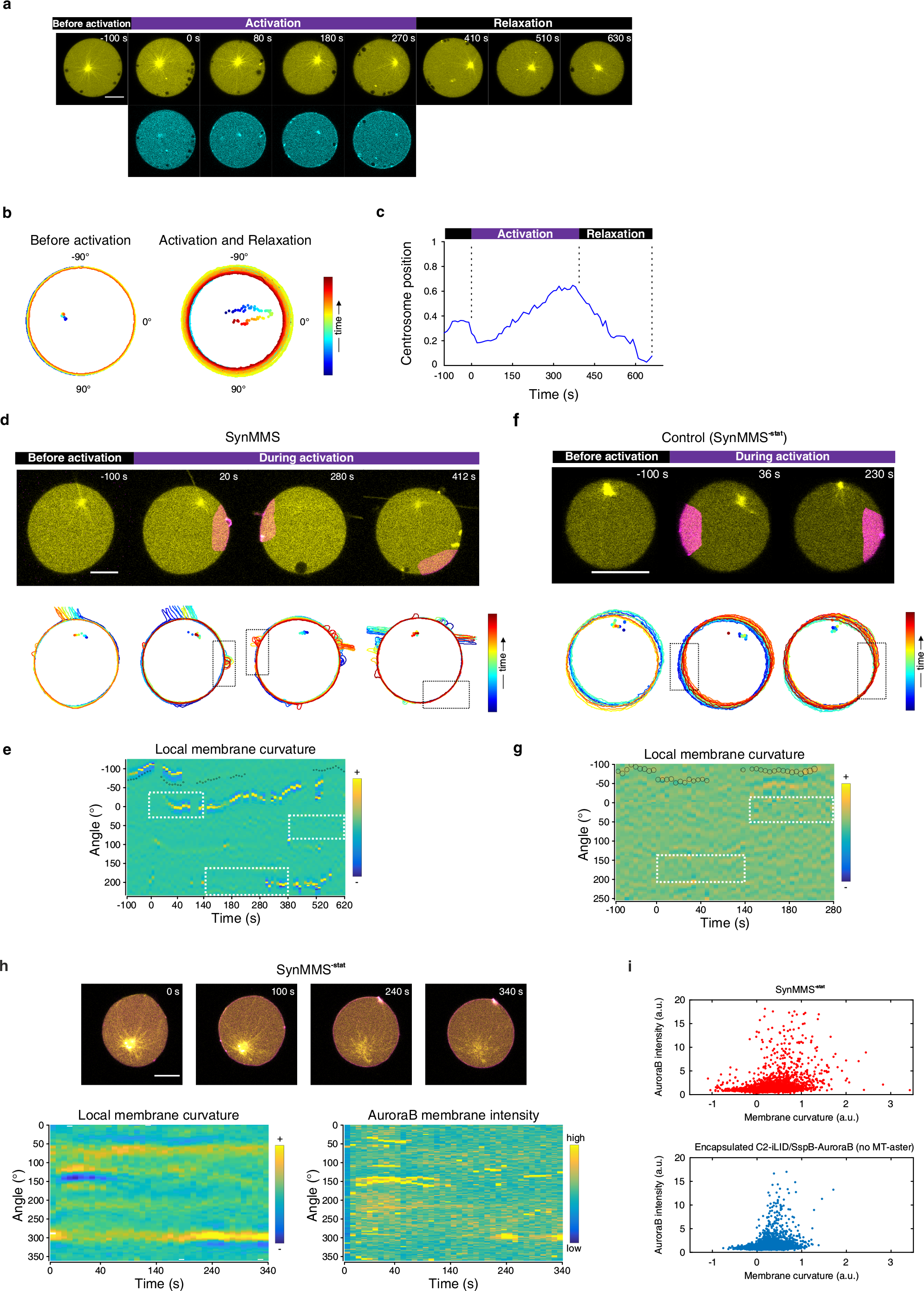
Light-induced AuroraB translocation generates a positive feedback between MT-growth and signaling at the deformable membrane. (a) Time lapse of Alexa647-tubulin (yellow, top) and Alexa488-SspB-AuroraB (cyan, bottom) in a GUV with encapsulated MT-aster/signaling system and a rigid membrane obtained before, during, and after global illumination with blue light. (b) Corresponding GUV contours for the entire time-lapse, color coded by time progression (dots: corresponding centrosome positions). Corresponding time evolution of centrosome position. 0 – centered, 1 – membrane proximal. Top row: Time lapse of Alexa647-tubulin (yellow) overlaid with Alexa488-SspB-AuroraB (magenta) at the equatorial plane of a SynMMS sequentially activated in different areas. Bottom row: GUV contours corresponding to above phases. Black rectangles mark activation regions. See also Supplementary Fig. 6a-f. (e) Kymograph of local membrane curvature (right: color code) corresponding to (d) overlaid with centrosome position (small circle: central, large circle: peripheral). White dashed rectangles denote the activation regions over time. (f) Time lapse and GUV contours as in (d) for a control SynMMS^−**stat**^ (without stathmin) sequentially activated at opposite sites. (g) Kymograph of local membrane curvature corresponding to (f). (h) Top row: time lapse of Alexa647-tubulin (yellow) overlaid with Alexa488-SspB-AuroraB (magenta) at the equatorial plane of a SynMMS^−**stat**^. Bottom: corresponding kymographs of local membrane curvature (left) and Alexa488-SspB-AuroraB membrane intensity (right). (i) Scatter plots of local membrane curvature versus Alexa488-SspB-AuroraB membrane intensity derived from (top) a set of 3 SynMMS^−**stat**^ (as in Supplementary Fig. 6d) and (bottom) a set of 5 GUVs with encapsulated C2-iLID/SspB-AuroraB and no MT-aster. Scale bars 10 μm.

To explore how kinase translocation affects this phosphorylation cycle and thereby the tubulin sequestration abilities of stathmin, we simulated the interaction between tubulin and stathmin with a reaction-diffusion model using measured enzymatic and association/dissociation parameters (Supplementary Fig. 4d,e; Supplementary Table 1; Methods). This showed that recruiting SspB-AuroraB activity to the membrane can locally overcome the luminal PPλ activity, yielding a pStathmin gradient emanating from the membrane (Fig. 3g). The numerical simulations also revealed that this gradient coincides with a gradient of free tubulin (Fig. 3g; Supplementary Fig. 4f,g), showing that the stathmin phosphorylation cycle constitutes an ATP-driven molecular ‘pump’ that maintains an out-of-equilibrium concentration of tubulin near the membrane (Fig. 3h).

To measure and characterize the predicted signal-induced gradient, we encapsulated COPY° together with the signaling system in GUVs. Ratiometric FRET measurements showed that COPY° was maintained in a dephosphorylated steady-state in the lumen in the presence of phosphatase activity (Supplementary Fig. 5a), whereas light-induced translocation of SspB-AuroraB resulted in steep (~0.5 μm) inward gradients from the GUV membrane (Fig. 3i,j; Supplementary Fig. 5b-e). This gradient only occurred in the presence of PPλ, showing that it was dynamically maintained by a phosphorylation cycle. Strikingly, these gradients mainly emanated from patterned structures, most likely originating from the spatially organized clusters of active SspB-AuroraB (Fig. 3i; Supplementary Fig. 5f). This signaling system thus embodies cytoplasmic reaction cycles in cells that maintain a low level of phosphorylated stathmin in the absence of stimuli, before a signal-induced recruitment of a kinase changes this kinetic balance and thereby locally maintains an enhanced concentration of tubulin near the membrane.

**Fig. 5.**
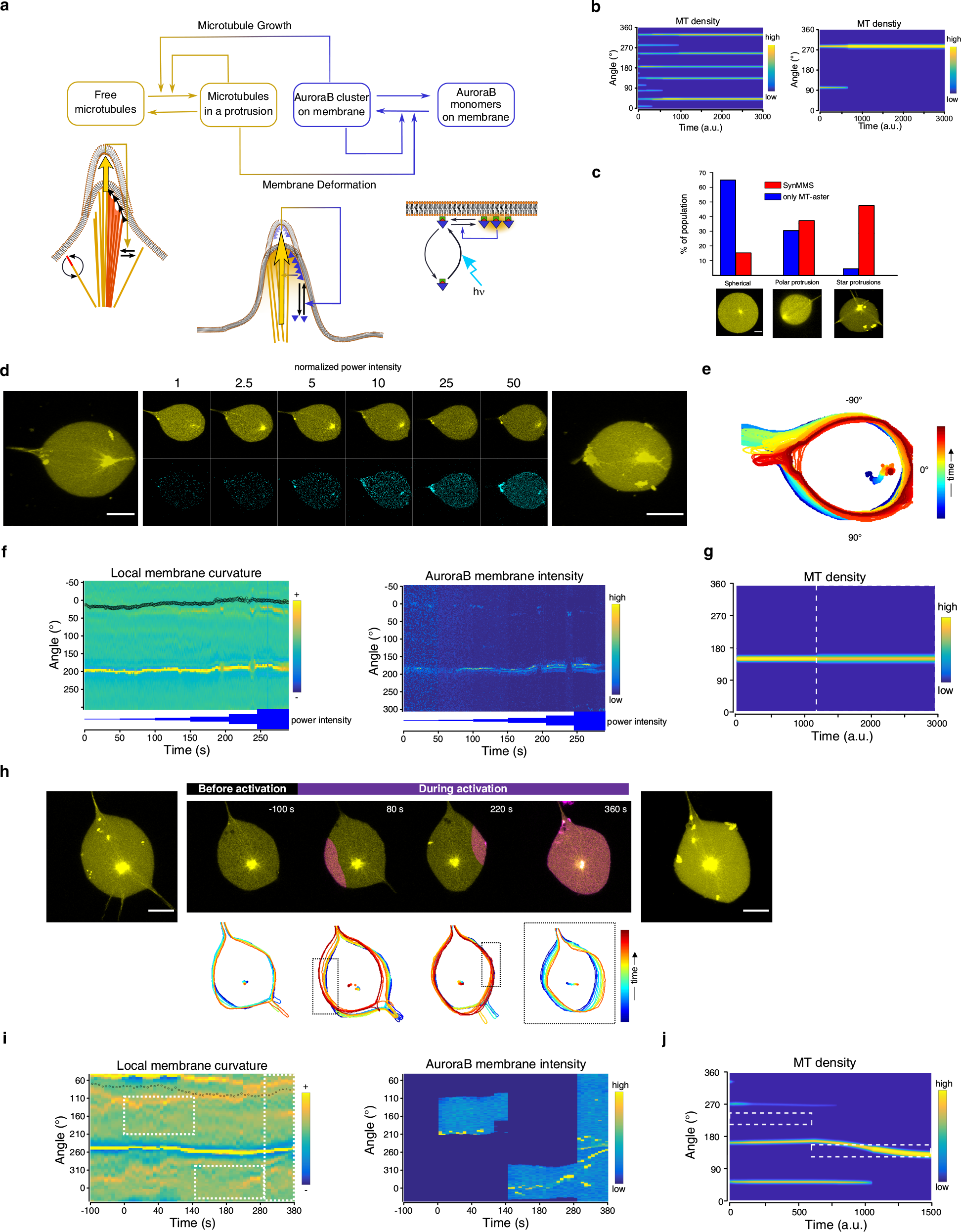
Reaction-diffusion model of the joint MT-aster/signaling dynamical systems predict SynMMS morphology distribution and stability. (a) The MT-aster/membrane (lower left, as in Fig. 2e) and AuroraB sub-systems (lower right, as in Fig. 3e), can be individually described by self-amplification of local structures (AuroraB clusters or MT-protrusions, feedback arrows) that trigger depletion of its free substrate (AuroraB monomers or free MTs, horizontal arrows). In SynMMS, a joint dynamical system is established: enhanced AuroraB clustering leads to MT-growth, which promotes MT-protrusions by capture of free MTs. This generates further membrane deformation which further enhances AuroraB clustering, resulting in a positive feedback. Membrane-proximal tubulin gradient depicted by yellow shading. (b) Simulated kymographs of MT density describing the evolution of the joint dynamical system. Left: when the rate of self-amplified AuroraB clustering on the membrane is greater than the local amplification rate of MT capture in SIC protrusions, the system evolves to a star-like morphology. Right: for the inversed case where self-amplified capture dominates over AuroraB self-amplification it evolves to a polar morphology. Parameters: see Methods. (c) Classification of experimentally obtained SynMMS morphologies (red) and encapsulated MT-asters (blue), in the absence of stimulus. Representative images of the three main morphological classes shown below. (d) Time-lapse of Alexa647-tubulin (yellow) and Alexa488-SspB-AuroraB (cyan) at the equatorial plane of a polar SynMMS during step-wise increase of global illumination with blue light. Maximum intensity projections of Alexa647-tubulin fluorescence of a confocal z-stack obtained before (left) and after (right) the dose-response series are shown. Light power intensity normalized to 1.8 mW/cm^2^. (e) Corresponding GUV contours for the entire time-lapse, color coded by time progression (dots: corresponding centrosome positions). (f) Kymographs of local membrane curvature (left; overlaid with centrosome position, small circle: central, large circle: peripheral) and SspB-AuroraB intensity along the membrane (right). Light power increase indicated by thickness of blue line, SspB-AuroraB intensity was normalized to the total intensity per frame. (g) Numerically obtained kymograph of MT density depicting the temporal evolution of a polar initial morphology subjected to global external signal (denoted by white dashed box). (h) Top row: Time lapse of Alexa647-tubulin (yellow) overlaid with Alexa488-SspB-AuroraB (magenta) at the equatorial plane of a polar SynMMS with a dominant protrusion sequentially activated on opposite sites followed by global irradiation. Bottom row: GUV contours corresponding to above phases of the time lapse. Black rectangles mark activation regions. (i) Corresponding kymographs of local membrane curvature and AuroraB intensity along the membrane for (h). White dashed rectangles denote the activation regions over time. Negative time: prior to activation. (j) Numerically obtained kymograph of MT density depicting the temporal evolution of a polar initial morphology with a dominant protrusion upon sequential exposure to localized external signals (denoted by white dashed boxes). The simulation demonstrates broadening of the strong protrusion, accompanied by the loss of minor protrusions. Scale bars 10 μm.

### A light responsive morphogenic vesicle system

To test if the light-induced membrane proximal tubulin gradient affects MT-growth, we encapsulated the signaling system together with MT-asters in GUVs with a rigid membrane. Light-induced SspB-AuroraB translocation resulted in centrosome decentering due to MT-growth (Fig. 4a-c; Supplementary Video 3), similarly to the temperature-induced MT-growth in a rigid GUV (Fig. 2c). Strikingly, the centrosome reverted to a central positioning within the GUV upon light removal (Fig. 4c), which shows that MT-growth can be reversibly activated by the reconstituted tubulin gradient-generating signaling system.

We next investigated if MT-induced membrane deformation can be brought about by activating the signaling system in GUVs with low membrane tension, a system to which we refer as a synthetic morphogenic membrane system (SynMMS). We transiently irradiated in sequence several SynMMS areas and monitored the formation and stability of the protrusions (Fig. 4d,e; Supplementary Video 5). Local irradiation on the right side immediately triggered the formation of a MT-induced protrusion, which persisted and evolved into a long protrusion after switching off the irradiation. Subsequent irradiation on the opposite side also induced the formation of persistent protrusions that remained after switching the irradiated area to the lower part. In this third area, no protrusions were formed, indicating that MTs were depleted by the protrusions formed during previous irradiations. In contrast, a control without stathmin (SynMMS^−**stat**^) did not form stabilized protrusions in any of the sequentially irradiated areas (Fig. 4f,g), which shows that stathmin is essential to couple the SspB-AuroraB signaling system to MT-dynamics (Supplementary Fig. 6a-g). SynMMS is thus capable of responding with directed and persistent morphology changes to local light cues.

**Fig. 6.**
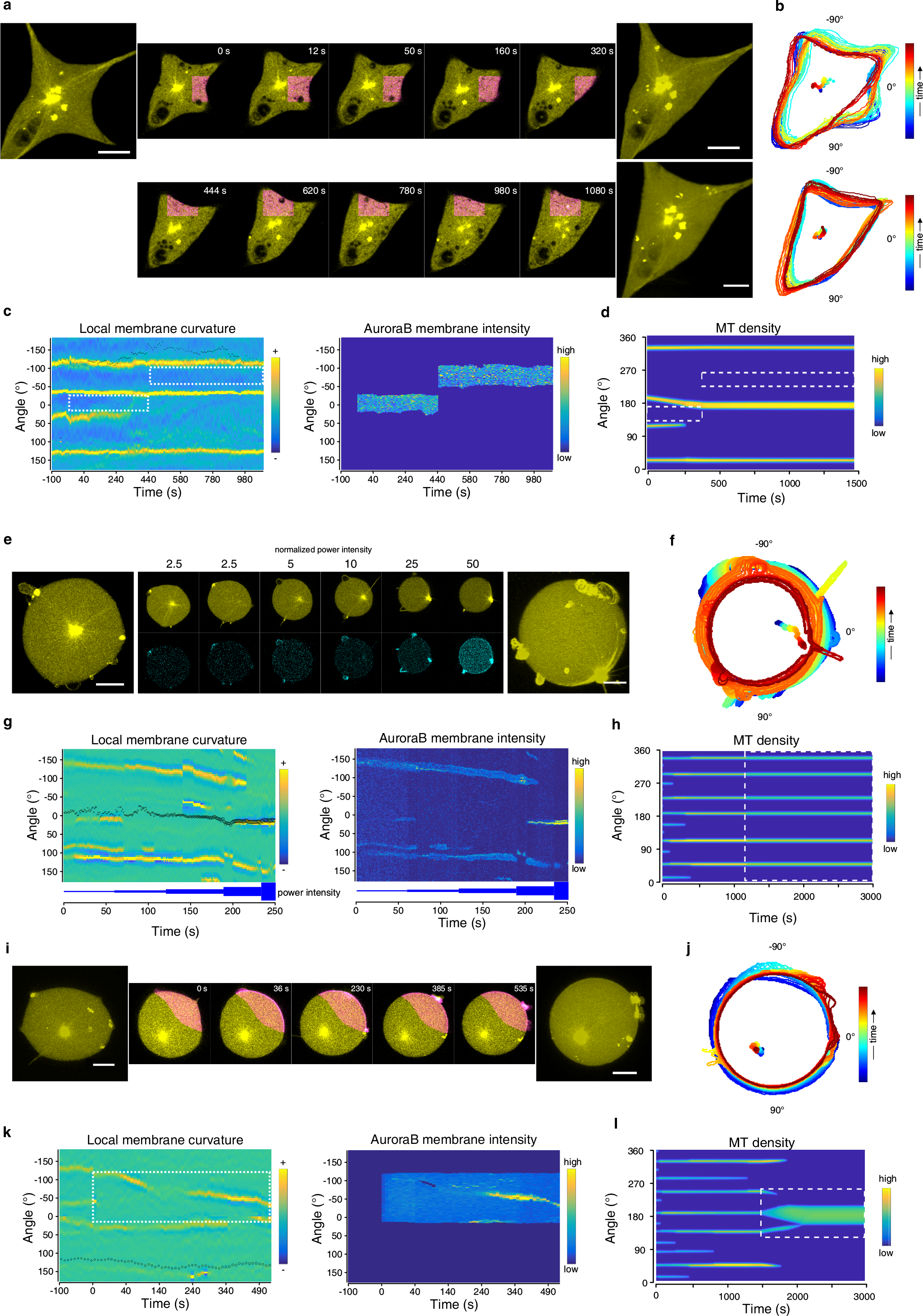
Global and local activation of star-like SynMMS induces morphological reorganization. (a) Top row: Time lapse of Alexa647-tubulin (yellow) overlaid with Alexa488-SspB-AuroraB (magenta) for a SynMMS with four strong SIC protrusions under sequential local light illumination (magenta area) between two protrusions. Bottom row: continuation of time lapse with local illumination in another region between two protrusions (magenta). Maximum intensity projections of Alexa647-tubulin fluorescence of a confocal z-stack obtained before (left) and after local irradiation (right) are shown. (b) Corresponding GUV contours for the entire time-lapse, color coded by time progression (dots: corresponding centrosome positions). (c) Kymograph of local membrane curvature (left; overlaid with centrosome position, small circle: central, large circle: peripheral) and SspB-AuroraB intensity along the membrane (right). White dashed rectangles denote the activation regions over time. (d) Numerical MT density kymograph indicating limits of plasticity of a star-like morphology stabilized by strong protrusions to subsequent spatially separated signals. White dashed rectangles denote the activation regions over time. (e) Time-lapse of Alexa647-tubulin (yellow) and Alexa488-SspB-AuroraB (cyan) at the equatorial plane of a star-like SynMMS during step-wise increase of global illumination with blue light. Maximum intensity projections of Alexa647-tubulin fluorescence of a confocal z-stack obtained before (left) and after (right) the dose-response series are shown. Light power normalized to 1.8 mW/cm^2^. (f) Corresponding GUV contours for the entire time-lapse, color coded by time progression (dots: corresponding centrosome positions). (g) Kymographs of local membrane curvature and SspB-AuroraB intensity as above. Numerically obtained kymograph of MT density depicting the temporal evolution of a star-like initial morphology subjected to a global external signal (denoted by white dashed box). (i) Time lapse of Alexa647-tubulin (yellow) overlaid with Alexa488-SspB-AuroraB (magenta) of a star-like SynMMS during continuous illumination in a localized region (magenta area) with blue light. Maximum intensity projections of Alexa647-tubulin fluorescence of a confocal z-stack obtained before (left) and after (right) the time lapse series are shown. (j) Corresponding GUV contours and (k) local membrane curvature kymograph and SspB-AuroraB intensity as above for (i). White dashed rectangles denote the activation region over time. (l) Numerically obtained kymograph of MT density depicting the evolution of a star-like morphology during exposure to a local signal (denoted by dashed white box). Local signal induces a star-to-polar morphological transition, through the coalescence of protrusions in the activated region. Scale bars are 10 μm.

The persistence of the protrusions after removal of the stimuli indicated that a feedback mechanism was responsible for their stabilization. To investigate if this signaling feedback was due to enhanced recruitment of SspB-AuroraB in MT-induced membrane deformations, we induced the translocation of SspB-AuroraB in a SynMMS^−**stat**^ (without stathmin) with a stable polar protrusion in which signaling cannot affect MT growth (Fig. 4h). Enhanced SspB-AuroraB was observed in the both transient and stable protrusions, consistent with the correlation between MT-induced membrane curvature and SspB-AuroraB intensity (Fig. 4i). This shows that SspB-AuroraB is preferentially recruited to MT-induced deformed membrane surfaces, likely caused by the local increased ratio of membrane surface to lumen volume in concave membrane deformations^10^, exhibiting enhanced signaling by a stronger tubulin gradient (Supplementary Fig. 6h). These experiments thus demonstrate that the deformable membrane establishes a positive feedback between signaling and cytoskeletal growth, which can sustain protrusions after stimuli removal, thereby enabling a morphological response that is dependent on stimuli history.

### Dynamical features of SynMMS enables self-organized morphogenic responses

To better understand how morphological behavior emerges from the interactions between the cytoskeletal and the signaling system, we conceptualized the dynamics of this coupled system with paradigmatic reaction-diffusion models. The behavior of each interacting sub-system can be described through self-amplification of local structures (AuroraB clusters or MT-protrusions) that trigger depletion of its free substrate (AuroraB monomers or free MTs) (Fig. 5a). Both sub-systems can independently break symmetry, reflecting spontaneous polar GUV morphology (Fig. 2), or self-organized SspB-AuroraB patterning (Fig. 3c-e). However, when interacting as in SynMMS, a joint dynamical system is established, where the MT-induced membrane deformations enhance SspB-AuroraB translocation and thereby its activity, that in turn locally promotes MT-growth (Fig. 5a). Reaction-diffusion simulations of this coupled system demonstrated that a new star-like morphology emerges for the joint system when the rate of self-amplified AuroraB clustering on the membrane is greater than the local amplification rate of free MTs in the SIC protrusions (Fig. 5b,left). In contrast, for the same total concentrations but inversed strength of the auto-amplification rates, the system evolves towards a single stable SIC protrusion, analogous to the dynamics of the cytoskeletal sub-system (Fig. 5b,right). This implies that the basal activity of the signaling system in absence of a stimulus accounts for the observed differences in initial morphologies between SynMMS and the encapsulated cytoskeletal system alone. This is also reflected in the bifurcation analysis of the coupled system which shows that the SspB-AuroraB concentration on the membrane determines if and how the system can break symmetry and thereby acquires self-organized shapes (Supplementary Fig. 7a). Indeed, quantification of the SynMMS morphologies using two morphometric classification parameters (Supplementary Fig. 7b, Methods) identified three coarse-grained morphological classes - spherical (semi-asters), polar, and star-like - whereas for the encapsulated cytoskeletal system alone, spherical and polar morphologies comprised the majority of shapes.

We next investigated both theoretically and experimentally the limits of morphological plasticity to global or local extracellular signals as a function of the initial shape. We speculated that strong protrusions would confer stability to the morphology, and therefore we first globally irradiated a polar SynMMS by step-wise increase of the light intensity (Fig. 5d-f; Supplementary Video 6). SspB-AuroraB was preferentially recruited to the pre-existing polar protrusion, which led to a light-dose dependent broadening of the main protrusion, indicating that enhanced signaling from deformed membrane areas in turn increases MT-growth. This was in agreement with simulations of the joint dynamical system where global stimulation only widened and further stabilized the pre-existing polar protrusion (Fig. 5g). Similarly, local stimulation of the regions flanking the main protrusion of a polar SynMMS also resulted in the polar protrusion reinforcement (Fig. 5h,i; Supplementary Video 6). The SynMMS rotated in the direction of the light signal, showing that overall MT-growth was indeed promoted towards the subsequently irradiated areas. However, during this reorganization, the sliding of MTs into the main protrusion could be observed which was accompanied by the clear loss of the lower three protrusions. Numerical simulations describing the effect of sequential flanking activation on a similar pre-patterned morphology, were in agreement with these observations (Fig. 5j). This demonstrates that a SynMMS, which acquired polar morphology through spontaneous symmetry breaking, will reinforce its polar state upon global as well as local light irradiation.

As already shown numerically, shape stabilization of the joint system can be either dominated by self-amplified MT-capture sites (Fig. 5a, left), or by the self-amplification of signaling (Fig. 5a, right). We thus further investigated the morphogenic plasticity of a star-like SynMMS whose shape was stabilized by four strong protrusions, indicative of dominating amplification of MT capture in SIC protrusions over signaling (Fig. 6a-c; Supplementary Video 7). Local irradiation between two protrusions of different size caused the coalescence of the smaller into the larger one, while subsequent irradiation between the newly enhanced and the neighboring large SIC protrusion did not trigger their coalescence. This shows that SIC protrusions in fact exchange MTs and a local tubulin gradient can bias capture towards a stronger protrusion, where large protrusions are stabilized due to a slow escape rate of the MTs, rendering them robust to local light stimuli in agreement with numerical findings based on a similar initial morphology (Fig. 6d).

On the other hand, our theoretical findings suggested that star-like initial states that are dominated by signaling (small protrusions) would exhibit the highest morphogenic plasticity. We therefore locally and globally stimulated star-like SynMMS with initially isotropically distributed small protrusions with few MTs, indicative of strong basal SspB-AuroraB activity. Increasing the global light-dose also enhanced growth of MTs in pre-existing protrusions, (Fig. 6e-g; Supplementary Video 7) which again shows that enhanced signaling in pre-formed membrane protrusions stabilizes them, in accordance with the numerical results (Fig. 6h). However, in contrast to a polar morphology, the growth of MTs resulted in strong centrosome decentering, promoting a transition to a spherical morphology with cortical MTs. We observed such decentering behavior in a set of SynMMS, but not in SynMMS^−**stat**^(Supplementary Fig. 7c). This suggested that morphologies with multiple protrusions that are stabilized by signaling can undergo global morphological transitions.

We thus asked if a local signal would result in an overall morphological transition towards the cue. In fact, upon local continuous illumination of a star-like SynMMS with multiple small protrusions, pre-existing membrane protrusions converged into the illuminated area, as theoretically predicted (Fig. 6i-l; Supplementary Video 7). Furthermore, as the protrusions migrated into this area with high SspB-AuroraB activity, MT-growth was further amplified leading to sprouting of additional micro-protrusions, as also observed in other SynMMS with strong signaling (Supplementary Fig. 6g). This resulted in an overall morphing process that went from star-like into a polar morphology and reorientation towards the irradiated region, demonstrating star-like SynMMS are capable of globally reorganizing their morphology in the direction of a localized stimulus.

These results thereby show that a localized stimulus can evoke a global morphological reorganization in SynMMS. However, the plasticity of this process is dictated by the prior morphological state. In an initial morphology that is stabilized by basal signaling activity (star-like), the accelerated growth of few MTs is sufficient for the maintenance of small protrusions. This leaves enough free MTs to continuously reorient existing protrusions in the direction of external signals to be eventually locked in by strong amplification of signaling. In contrast, dominant SIC protrusions constrain this morphological plasticity by acting as strong MT sinks. Thus, signaling strength at the plasma membrane confers plasticity to morphological responses, whereas strong SIC protrusions constrain the morphological transitions.

## DISCUSSION

To distill basic principles of morphogen-guided cell morphogenesis, we reconstituted dynamic microtubule asters inside GUVs together with a signaling system that is actuated by dimensionality reduction and operates under non-equilibrium conditions. We found that light-induced translocation of SspB-AuroraB induces a stathmin phosphorylation cycle near the membrane, thereby enhancing the local tubulin concentration which promotes MT growth. Such a tubulin ‘pump’ could serve as a generic mechanism for directed MT-growth, e.g. guiding MTs to kinetochores during pro-metaphase in mitosis, where AuroraB localized on centromeres drives stathmin phosphorylation^32,34^.

This coupling between signaling and MT-dynamics is however not uni-directional. We identified that the reverse causality is also established, enabling spontaneous morphogenesis to occur, and our findings identified the deformable membrane as a key mediator of this positive feedback mechanism. This implies that it occurs in mammalian cells that generate shapes by deforming the membrane with the MT-cytoskeleton. We could demonstrate that this system without direct molecular feedback already has self-organized morphogenic abilities based on the principle of local matter coalescence through self-amplification, which in turn depletes the substrate from the surrounding. This is analogous to Grassé’s stigmergic concept of collective behavior of social insects through indirect communication that lead to self-organized creation of nest structures^35^. The biomimetic design of SynMMS to respond to localized light cues with a translocation gradient of SspB-AuroraB that mimics the response to an extracellular morphogen gradient, revealed that directional morphological responses to extracellular templates arise from constraining intrinsic self-organized dynamical solutions of stigmergic systems^36^.

Although many layers of biochemical regulation likely contribute to both morphological plasticity and stabilization in cells^37,38^, we demonstrate that the recursive interaction of MTs with signaling at the deformable membrane is sufficient to fulfill both requirements. Our observations that the star-morphology is the most plastic, whereas the polar the most robust one, lead to the question how cells can switch between plasticity and stabilization of shapes. In this sense, the interplay between actin and MTs may be of fundamental importance, where actin would generate rapid and local morphological dynamics at the cell periphery, while the MT-cytoskeleton would guide the global shape. This could be further elucidated using synthetic systems that build upon SynMMS from the progress made in reconstituting actin networks on and within synthetic membranes^39,40^. An additional step would be the inclusion of MT-interacting motor proteins that can further affect the self-organization of MTs by regulating MT dynamics and bundling^15,41,42^.

At the higher scale, steps have been made to generate synthetic tissues where porous proto-cells communicate by diffusive protein signals to form larger tissue-mimicking arrays^43^. The communication in these synthetic systems was based on distinct signal emitter and receiver synthetic cells. The intrinsic self-organizing morphological dynamics of SynMMS has the potential to establish recursive communication between SynMMS^44^, enabling the investigation of basic principles of self-organized tissue formation.

The short life span of SynMMS is constrained by the encapsulated fuel store (ATP, GTP). By equipping SynMMS with an artificial photosynthetic membrane^45^, light energy could be converted into ATP or even GTP chemical potential^46^ that could maintain its non-equilibrium state over prolonged periods. Finally, recent progress in reconstituting self-replicating DNA by its encoded proteins in liposomes^47^, bacterial cell division^12^ and lipid metabolism^48^ suggests that self-replicating properties could be conferred to synthetic systems such as SynMMS in the future.

## ACKNOWLEDGMENTS

The authors thank Peter Bieling for reagents, helpful discussions and help in shaping the manuscript, Roger Goody for assistance with the stopped-flow experiments, H. Schütz, K. Michel, M. Reichl, and S. Gentz for technical assistance and A. Krämer for critical reading of the manuscript. This project was funded by the BMBF/MPG network MaxSynBio (031A359A).

## AUTHOR CONTRIBUTIONS

K.G. developed biochemical assays as well as COPY°, purified and encapsulated proteins, performed biochemical and imaging experiments. F.G. generated encapsulated asters, performed MT assays and imaging experiments, B.S. developed and performed image and morphometric analysis, A.N. and M.C.M. performed numerical simulations, H.S. performed biochemical assays and imaging experiments, M.S. developed and performed gradient reaction-diffusion simulations, A.K. & P.B. developed theoretical concepts, P.B. conceived and supervised the project and wrote the manuscript with help of A.K., B.S. and K.G. All authors provided input to the preparation of the manuscript.

## METHODS

Detailed methods are provided in the online version of this paper and include the following:

- CONTACT FOR REAGENT AND RESOURCE SHARING
- METHOD DETAILS

○ Preparation of recombinant proteins
○ Preparation of tubulin and centrosomes
○ Protein labeling
○ Protein encapsulation in GUVs by cDICE
○ Imaging of MT asters and morphological states in GUVs
○ Single-filament TIRF-M assay and data analysis
○ Determination of MT-aster size in vitro
○ Morphometric analysis of GUVs
○ Imaging and analysis of light-induced translocation
○ Quantifying regularity of SspB-AuroraB clustering on the membrane
○ COPY° characterization
○ Stopped-flow measurements
○ Determination of enzymatic kinetic parameters
○ Measurement of stathmin phosphorylation gradients
○ Paradigmatic model of morphogenesis
○ Reaction-diffusion simulation of gradients

## SUPPLEMENTAL INFORMATION

Supplemental Information includes seven figures, two tables, seven videos and can be found with this article online

## METHODS

### CONTACT FOR REAGENT AND RESOURCE SHARING

Further information and requests for resources and reagents should be directed to and will be fulfilled by the corresponding author, Philippe Bastiaens.

## METHOD DETAILS

### Preparation of recombinant proteins

Amino acid sequences of all used constructs are presented in Supplementary Table 2. All proteins were expressed in *E. coli* BL21 DE3 RIL over night at 18 °C after induction with 0.2 mM IPTG. AuroraB was bicistronically expressed with the INCENP in-box from a modified pGEX6P-2rbs plasmid, which was a gift from A. Musacchio. All other proteins were expressed from a pET24 plasmid as fusions to an N-terminal His_10_-tag followed by a TEV protease site. 4 – 6 liters of LB medium were used to express Stathmin, iLID, and λ-phosphatase (λ-PPase), whereas 10 – 12 liters of TB medium were used for AuroraB. Cells were lysed by two passes through an Emulsiflex C5 (Avestin, Mannheim, Germany), and the lysate was cleared by 45 – 60 min centrifugation at 20000 rpm in an A27-8×50 rotor (Sorvall/Thermo Fisher Scientific, Dreieich, Germany).

#### Stathmin and iLID

Gly-Stathmin-Cys, Gly-iLID-tRac1, or Gly-C2-iLID were purified via Ni-NTA Superflow (Qiagen, Hilden, Germany) in 50 mM NaP_i_ pH 8, 300 mM NaCl, 2 mM MgCl_2_, 0.1 mM β-mercaptoethanol, 10 mM imidazole. The column was washed with 60 mM imidazole, and the protein eluted by a step to 500 mM imidazole. The eluate was supplemented with 2 – 4 mg TEV protease and dialyzed over night against 2 L of 50 mM Tris pH 8, 100 mM NaCl, 1 mM EDTA, 0.1 mM β-mercaptoethanol. The cleaved protein was adjusted to 10 mM imidazole (60 mM for Gly-C2-iLID), and passed again through the Ni-NTA column, followed by gel filtration on a HiLoad 26/600 Superdex 75 pg column (GE Healthcare, Solingen, Germany) in 50 mM Hepes pH 7.5, 200 mM NaCl, 2 mM MgCl_2_, 2 mM DTT. For iLID constructs, the sample was adjusted to a saturated concentration of flavin mononucleotide prior to gel filtration. Gly-iLID-tRac1 was subsequently geranylgeranylated as described elsewhere ^49^. We refer to geranylgeranylated iLID as iLID_G.

#### λ-phosphatase

The protein was purified via cobalt-loaded HiTrap Chelating HP columns (GE Healthcare) in 50 mM NaP_i_ pH 8, 500 mM NaCl, 2 mM MgCl_2_, 5 % v/v glycerol, 0.1 mM β-mercaptoethanol, 10 mM imidazole. The washing/elution steps were 60, 200, 500 mM imidazole. The eluted protein was quickly supplemented with 10 mM DTT and digested with TEV protease as above, while dialyzing against 2 L of 50 mM Tris pH 8, 500 mM NaCl, 5 % v/v glycerol, 2 mM MgCl_2_, 10 mM DTT. The cleaved protein was buffer exchanged back into the sodium phosphate buffer via a HiPrep Desalting 26/10 column (GE Healthcare), adjusted to 10 mM imidazole, and passed again over the chelating column. Finally, the protein was gel filtered on a HiLoad 26/600 Superdex 75 pg column (GE Healthcare) in 50 mM Hepes pH 7.5, 500 mM NaCl, 2 mM MgCl_2_, 5 mM DTT, 10 % v/v glycerol.

#### AuroraB

Gly-SspB-AuroraB(45-344)/INCENP(834-902) was purified via Glutathione Sepharose 4 Fast Flow (GE Healthcare) in 50 mM Hepes pH 7.3, 500 mM NaCl, 2 mM MgCl_2_, 5 mM DTT, 5 % v/v glycerol. The column with bound protein was washed with buffer containing 1 mM ATP, then the protein was eluted with buffer containing 20 mM glutathione and additional DTT up to 10 mM. The protein was cleaved over night with TEV protease while remaining in the elution buffer. The cleaved protein was buffer exchanged via a HiPrep Desalting 26/10 column (GE Healthcare) to remove glutathione, and passed again through the Glutathione Sepharose column. Gel filtration was carried out on a HiLoad 26/600 Superdex 200 pg column (GE Healthcare) in 50 mM Hepes pH 7.3, 500 mM NaCl, 2 mM MgCl_2_, 5 mM DTT avoiding concentrating the protein before gel filtration. Finally, the protein was concentrated in an Amicon Stirred Cell (Merck Millipore, Darmstadt, Germany), adjusted to 20% glycerol and hard spun at 400.000 g prior to shock freezing.

### Preparation of tubulin and centrosomes

Tubulin purification was performed according to a standard method ^50^. The following deviations from the published procedure were made: the ratio of buffer/brain mass was 1.5 l/kg, and the Pipes final concentration in the polymerization steps was 333 mM. Purified tubulin was adjusted to a concentration of 200 μM with BRB80 (80 mM Pipes pH 6.8, 1 mM MgCl_2_, 1 mM EGTA) and stored at −150 °C.

Tubulin was labeled with NHS-Alexa568, −488 or −647 (Thermo Fisher Scientific) or EZ-link NHS-biotin (Thermo Fisher Scientific) according to a standard procedure^51^. Typically, 50 mg of tubulin were labeled with a 7.5-fold molar excess of dye assuming 70 % tubulin recovery after the initial polymerization step. Following deviations from the published procedure were made: the dye was added in two steps, each followed by a 15 min incubation, totaling 40 μl DMSO in 1 ml buffer. The reaction was not specifically stopped, and all ultracentrifugation steps were performed at an increased speed of 400.000 g in an MLA-80 rotor (Beckman Coulter, Krefeld, Germany). Labeled tubulin was adjusted to 200 μM with BRB80 and stored at –150 °C.

Centrosomes were isolated from KE37 cells following a standard method ^52^.

### Protein labeling

Proteins were specifically labeled by sortagging ^53^. The LPETGG peptide was conjugated to NHS-Alexa488, −647, or Atto532 over night at 30 °C (20 mM peptide, 40 mM dye (Thermo Fisher Scientific for Alexa dyes, Atto-tec (Siegen, Germany) for Atto532) in DMSO), the reaction stopped by 100 mM Tris pH 8. Typically, 1 – 3 mg protein (ideally around 300 μM final concentration in the reaction), were mixed with a 4-fold excess of conjugated peptide, ca. 100 μM Sortase A (from *S. aureus*, gift from P. Bieling), and 6 mM CaCl_2_. For iLID constructs, the sample was additionally adjusted to a saturated concentration of flavin mononucleotide. The reaction was allowed to proceed over night at 18 °C or 4 °C (for λ-PPase and AuroraB), and the mixture was separated by gel filtration on a Superdex 75 10/300 GL column (GE Healthcare) in appropriate protein gel filtration buffer.

### Protein encapsulation in GUVs by cDICE

Encapsulation of proteins in GUVs was achieved by continuous Droplet Interface Crossing Encapsulation following the original method ^28^ with relevant parameters noted below. Empty GUVs used for experiments in the outside configuration were also produced by this method.

0.36 mM lipids were prepared in mineral oil (M3516, Sigma Aldrich/Merck, Darmstadt, Germany). The mixture consisted of eggPC (Sigma Aldrich/Merck) with 15 mol% DOPS and 0.05 mol% DOPE-biotin (both from Avanti Polar Lipids, Alabaster, U.S.A.). Additionally, 0.05 ml% LissamineRhodamineB-DOPE was added for experiments to analyze Alexa488-SspB-AuroraB or Alexa647-SspB-PPλ patterning on GUV membranes. Glass capillaries were typically 10 – 15 μm wide, and pressure was applied using an MFCS-EZ control system (Fluigent, Jena, Germany). cDICE chambers were 3.5 cm wide and were rotated at 1500 – 1800 rpm. 8-well Lab-Tek chambers (Thermo Fisher Scientific) or 18-well μ-slides (ibidi, Martinsried, Germany) were coated with streptavidin for GUV immobilization.

The base cDICE buffer contained 80 mM Pipes pH 6.8, 75 mM KCl, 2 mM MgCl_2_, 1 mM EGTA, 1 mM trolox, 0.1 mg/ml β-casein. Additionally, the inner buffer contained 300 mM sucrose and 80 mM glucose, while the outer buffer contained 380 mM glucose. Whenever an osmotic deflation of GUVs to decrease membrane tension was desired, the sucrose concentration inside was lowered by 60 mM and the outside glucose concentration was increased by 160 mM (i.e. Fig. 2a,e,d; Figs. 4,5,6; Supplementary Fig. 1g, Supplementary Fig. 6a,d). To induce deflation of GUVs for Supplementary Fig. 1f, 50 μl of 1 M glucose were added to 300 μl of GUVs.

In experiments involving PPλ, EGTA was replaced by 0.8 mM MnCl_2_. For tubulin polymerization, 2 mM GTP was included, whereas phosphorylation gradient experiments just required the presence of soluble tubulin, for which 60 μM GTP were included. For AuroraB, 2 mM ATP was added, and in experiments requiring both nucleotides the MgCl_2_ concentration was increased to 4 mM. Oxygen scavengers (see “Single-filament TIRF microscopy and data analysis”) were included in cases with just microtubule asters and stathmin, but were omitted in all other encapsulation experiments.

The encapsulation efficiency of tubulin and stathmin was determined by a reference concentration outside of GUVs (Supplementary Fig. 2c), and the used protein concentration in cDICE was adjusted to yield desired final concentrations, as indicated in respective experiments. For encapsulation of centrosomes, the samples typically contained 10 vol% of a purified centrosome fraction.

For the encapsulation of iLID, AuroraB, and PPλ, an exact concentration was not targeted, but the resulting inside concentrations and membrane densities given specific starting concentrations were quantified (Supplementary Fig. 3c) and verified to result in sufficient membrane recruitment and desired stathmin phosphorylation state (see “Imaging and quantification of light-induced protein translocation” and “Measurement and analysis of local stathmin phosphorylation gradients”).

Prior to cDICE, the samples were centrifuged for 15 min at 100.000 rpm in a TLA-100 rotor (Beckman Coulter). The centrosomes were never centrifuged, while the tubulin stock was centrifuged separately for 30 min at 60.000 rpm in an RP80-AT2 rotor (Sorvall/Thermo Fisher Scientific).

### Imaging of MT asters and morphological states in GUVs

To produce MT asters in GUVs, soluble tubulin and centrosomes were encapsulated as described in “Protein encapsulation in GUVs by cDICE”. Stathmin was included as indicated for individual experiments. The following protein concentrations were used in the full reconstituted morphogenic system: 40 or 44 μM (final) tubulin (10 % Alexa647-tubulin), 4 μM (final) stathmin, 5 μM Gly-C2-iLID, 12 μM Alexa488-SspB-AuroraB/INCENP, 1 μM λ-PPase. Alexa568-tubulin was used for experiments without the signaling system at indicated concentrations. Alexa488-tubulin was used in experiments to establish dependency between membrane curvature and Alexa647-SspB-PPλ membrane recruitment.

Imaging was done on a Leica SP8 confocal microscope (Leica Microsystems, Wetzlar, Germany) equipped with Leica HyD hybrid detectors at 33 °C in an environment-controlled chamber (Life Imaging Services, Basel, Switzerland) using an HC PL APO 63×/1.2NA motCORR CS2 water objective (Leica Microsystems). The elevated temperature promoted tubulin polymerization, the asters reaching their end state in ca. 7 minutes, after which single-plane images or z-stacks (1 or 0.5 μm spacing) were taken. For the temperature ramp experiments, GUV-containing Lab-Tek chambers were incubated at 4 °C for 10 minutes in the pre-cooled Stable Z Lab-Tek Stage (Bioptechs, Butler, U.S.A.). Then, the stage was moved onto the microscope equilibrated at room temperature and reached RT (~ 21 °C) in 10 minutes. The measurements indicated as “cold” were done under this condition. After that the Stable Z Lab-Tek Stage controller was used to heat the sample to 34 °C within 20 minutes and images were taken every minute.

Alexa568 or Alexa488 were excited with a 470 – 670 nm white light laser (white light laser Kit WLL2, NKT Photonics, Köln, Germany) at 578 nm (270 – 600 mW/cm^2^) or 488 nm (220 – 340 mW/cm^2^), respectively. Detection of fluorescence emission was done in standard or counting mode, restricted with an Acousto-Optical Beam Splitter (AOBS) to 588 – 640 nm or 498 – 540 nm, respectively. Images were taken with 2-4× line averaging at 400 or 200 Hz scanning speed. The pinhole was set to 1 Airy unit here and in all experiments described in this work.

In the full morphogenic system, Alexa647 was excited at 650 nm (200 mW/cm^2^ - 700 mW/cm^2^) at 1000 Hz scanning speed to reduce photodamage. Detection of fluorescence emission was done in counting mode, restricted to 650 – 750 nm and further restricted by LightGate time-gating to minimize signal from laser reflection and background. Activation of iLID and imaging of Alexa488-SspB-AuroraB was done with 488 nm (220 mW/cm^2^) excitation from a white light laser source, restricting detection to 498 – 580 nm and by LightGate. Time series images and stacks (1 μm spacing) were taken with 8x line accumulation in line sequential scanning mode. For local activation, 488 nm illumination was restricted to a manually drawn ROI.

### Single-filament TIRF-M assay and data analysis

GMPCPP-stabilized microtubule seeds were prepared as follows: 40 μL of 40 μM tubulin mixture (25 % tubulin, 25 % Alexa568-tubulin, 50 % biotinylated tubulin) was polymerized in seed buffer (80 mM Pipes pH 6.8, 1 mM MgCl_2_, 1 mM EGTA, 1 mM GMPCPP (Jena Bioscience), 5 mM β-mercaptoethanol) at 37 °C. After 30 minutes, 400 μL of prewarmed BRB80 (80 mM Pipes pH 6.8, 1 mM MgCl_2_, 1 mM EGTA) was added to the tubulin mixture, which was then spun down in a table top centrifuge at 21.000 g for 8 minutes at RT. The microtubule pellet was thoroughly resuspended in 40 μL of BRB80.

TIRF flow chambers were assembled from a biotin-PEG functionalized glass coverslip attached to a PLL-PEG passivated cover slide using double-sided tape ^54^. To prevent potential unspecific binding, the flow chamber was first flushed and incubated with 35 μL of blocking buffer (100 β-casein μg/mL, 1 % w/v Pluronic F-127 in assay buffer (40 mM Pipes pH 6.8, 75 mM KCl, 4 mM MgCl_2_, 0.4 mM EGTA, 1 mM GTP, 100 β-casein μg/mL, 1 mM trolox, 20 mM β-mercaptoethanol). After 5 minutes, the flow chamber was washed two times with 35 μL of assay buffer. Then, the chamber was flushed and incubated with 35 μL of 150 μg/mL Neutravidin (Thermo Fisher Scientific) in assay buffer for another 5 minutes. The flow chamber was washed two times with 35 μL of assay buffer, and then the chamber was flushed and incubated with 35 μL of diluted (1:750) GMPCPP-stabilized microtubule seeds. After 5 minutes, the chamber was washed two times with 35 μL of assay buffer.

After the seeds were immobilized on the surface, 40 μL of protein mixture (10 – 40 μM tubulin (7.5 % Alexa568-labeled), 0 – 20 μM stathmin, 0 – 30 μM phospho-stathmin, 4 μL 10× oxygen-scavenging system ^54^, 0. 2% w/v methylcellulose) in assay buffer was introduced into the flow chamber and the two sides were sealed with transparent nail polish.

Imaging was performed at RT with a custom-built TIRF microscope (Olympus IX81 base) with a 60x Olympus APON TIRF objective with TOPTICA IBeam smart 560s laser and a Quad-Notch filter (400-410/488/561/631-640). Temperature was kept at 34 °C by a collar objective heater (Bioptechs). Image acquisition was done with an EM CCD Andor iXon 888 camera controlled by Micromanager 1.4 software. Fiji ImageJ was used for data analysis. For each condition, the average growth velocity and catastrophe frequency were determined from ≥60 microtubules by kymograph analysis.

### Determination of MT-aster size *in vitro*

20 μL of protein mixture (10 – 40 μM tubulin (7.5% Alexa568-labeled), 0 – 20 mM stathmin, 0 – 30 μM phospho-stathmin, 2 μl Silica beads (SS06N, Bangs Laboratories, Fishers, U.S.A.), 2 μl 10x oxygen-scavenging system^54^, 0.2 % w/v methylcellulose) was dispersed on a glass cover slide blocked with PLL-PEG^54^ and covered with a glass coverslip pre-coated with PLL-PEG. The edges were sealed with transparent nail polish. Imaging of asters at 33 °C or in temperature ramp experiments were done as described in “Imaging of microtubule asters and morphogenic state transitions in GUVs”. 12 μm thick z-stacks (1 μm spacing) were taken in the experiments at 33 °C, but single planes were taken during the temperature ramp to maximize the number of imaged fields of view.

For each individual condition, the z-stacks (if present) of each position were average projected in Z and aligned on each other, and projected again using a custom macro in ImageJ. The radial intensity profile of the averaged data (> 30 asters per condition) was extracted with a custom ImageJ macro. The aster size was determined as the 99 % decay length of a mono-exponential fit to the radial intensity profile in Origin (OriginLab, Northhampton, U.S.A.).

### Morphometric analysis of GUVs

#### Centrosome Position and Eccentricity

A mask of the GUVs was obtained by thresholding the fluorescence intensity of fluorescently labeled tubulin at the plane of the centrosome and an ellipse was fitted to the mask in ImageJ. The centrosome position relative to the center of mask was calculated by:

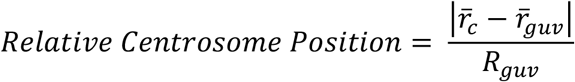

where 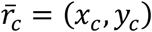 and 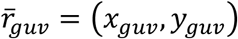 are the position of the centrosome and the geometric center of the fitted ellipse obtained in ImageJ, respectively, and *R*_*guv*_ = (*M* + *m*)/4 where *M* and *m* are the major and minor axis of the ellipse. The eccentricity of the fitted ellipse was calculated as:

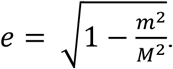

#### Morphometric Classification

A 3D mask of the GUV shape was made in the fluorescently labeled tubulin confocal Z-stack, by performing a 2-pixel Mean Filter and thresholding. Masks were imported to MATLAB 2018b. The centrosome position was calculated as above, except that in 3D, *R*_*guv*_ = (*PA*1 + *PA*2 + *PA*3)/6 where *PA*1,2,3 are the Principal Axis Lengths and 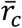 is the GUV solid mask centroid. These were obtained by applying the in-built function *regionprops3* to the GUV solid mask. The diameter D of the minimal bounding sphere was calculated using the *minboundsphere* function by John D’Errico available at the MathWorks File Exchange. The diameter of the minimal inbound sphere was calculated by taking the maximum of the distance transform of the points contained in the GUV mask, to the GUV surface (edge of the mask) by means of the in-built function *bwdist*.

#### Local curvature kymographs

At each time point of the time-lapse experiments, a mask of the GUV shape was made in the fluorescence image of labeled tubulin (except for Fig. 4h,i, where the mask was done on the labeled SspB-AuroraB image), by performing a 2-pixel Mean Filter and thresholding in Fiji. A spline was fitted to the Region of Interest (ROI) and then converted into a segmented line format. ROIs and CLSM images were imported to MATLAB 2018b. The points in the segmented line were ordered by their angle with respect to the geometric center of the GUV mask. The curvature was calculated with the function LineCurvature2D written by D. Kroon (University of Twente, August 2011) and available at the MathWorks File Exchange. The curvature at a point was calculated by considering as left and right neighboring points that were 5 points away from it.

#### Fluorescence intensity angular kymographs

For the Alexa488-SspB-AuroraB fluorescence intensity kymographs, the GUV mask at each time point was eroded and dilated using *imerode* and *imdilate* respectively, forming a band following the shape of the GUV. The mask was segmented into 230 angular bins, and the maximum intensity at each of them was taken.

#### Quantification of enhanced protein recruitment in protrusions

Local curvature as well as SspB-Aurora fluorescence intensity kymographs were made, where the masking was performed on the SspB-Aurora fluorescence image. The curvature at each kymograph pixel value was plotted against its corresponding pixel in the fluorescence kymograph.

### Imaging and analysis of light-induced translocation

Imaging of light-induced translocation of SspB-AuroraB (Alexa488- or Alexa647-labeled) or Alexa647-SspB-λ-PPase to membrane-bound iLID (Alexa488-labeled or unlabeled) was done on a Leica SP8 confocal microscope (see “Imaging of microtubule asters and morphogenic state transitions in GUVs”). iLID was activated by the 458 nm Argon laser line. The following laser powers were used to determine the dose-response properties of the iLID system: near maximal recruitment was observed at 30 mW/cm^2^, so that 65 mW/cm^2^ was used to observe recruitment kinetics and to activate the full morphogenic system. Alexa647 and Alexa488 were excited with 650 and 495 nm white line laser lines, respectively. Detection of fluorescence emission was done in photon counting mode, restricted with an Acousto-Optical Beam Splitter (AOBS) (505 – 560 nm for Alexa488, 660 – 700 nm for Alexa647), and further restricted by LightGate time-gating to minimize signal from laser reflection and background. Images were taken in sequential frame mode when using Alexa647-labeled proteins, where the 488 nm channel would be used to activate iLID and also image it in the cases when iLID was also labeled. For the translocation of Alexa488-labeled SspB-AuroraB, unlabeled iLID was used, and only images of the 488 nm channel were taken to simultaneously activate iLID and image SspB-AuroraB. Typically for activation, it was sufficient to take an image in the 488 nm channel every 3 – 4 s, but sometimes higher framerates (ca. 0.6 s) were used to fully resolve the kinetics.

For the outside experiments, varying protein concentrations were added to immobilized GUVs as indicated in individual Fig. legends. We mainly used C2-iLID, but we found no general differences in the behavior of the prenylated iLID_G in respect to recruitment and unbinding kinetics as well as to achievable surface densities of the effector proteins. The defined protein concentration in the bulk was used to calculate the protein densities on the membrane using a semi-automated tool in Matlab (MathWorks, Natick, U.S.A.) ^55^. Bulk depletion effects could be neglected because only 10 – 50 GUVs were present in each 50 μl sample. Binding and unbinding kinetics were exponentially fit in Origin (OriginLab).

For inside GUV experiments, defined starting concentrations are indicated in individual Fig. legends. To quantify the encapsulation efficiency and membrane densities, a defined protein concentration was added outside of the GUVs as a reference. Analysis was done in the same manner as for outside GUV experiments.

Local light-induced translocation resulted in a lateral gradient of membrane-bound proteins due to lateral diffusion, which was characterized as follows. A GUV encapsulating Alexa647-SspB-PPλ (SspB) and iLID was imaged for 140 s every 2 s, while irradiating a region of interest encompassing roughly 25% of the membrane for 25s (Supplementary Fig. 3e). The resulting redistribution of SspB in the GUV was fit by 2D CA-simulation with the following model: SspB has a minor affinity for the membrane as discernable by the difference in contrast in the images before photo-recruitment with a partitioning resulting from association and dissociation rate constants k_assoc_/k_dissoc_; irradiation “creates” a tightly membrane-bound fraction at a rate constant k_recruit_ that is released according to k_release_, but due to the amount of iLID encapsulated concentration of SspB at the membrane is capped at c_max_ (normalized intensity); both membrane-bound iLID and SspB can diffuse laterally (D_lat_) and SspB in solution diffuses with a coefficient D_sol_. The time-lapse of the experiment was normalized to account for bleaching and segmented to improve signal-to-noise. The simulation utilized a time resolution of 0.02 s and matched the pixel size of 0.181×0.181 μm of the confocal micrographs. By varying the parameters of the simulation via function lsqnonlin (Matlab), a best fit was achieved for D_lat_ = 26.4 μm^2^/s; k_recruit_ = 4.6/s; k_release_ = 0.016/s; k_assoc_/k_dissoc_ = 0.38; c_max_ = 4.2 a.u. The time-resolution of the experiment was too slow to resolve the very fast diffusion of SspB in solution, so any D_sol_ > 200 μm^2^/s fits the data well. Any discrepancy of kinetic values stems from the fact that the apparent binding and unbinding reaction fit in Fig. 3a is a convolution of diffusion, membrane association/dissociation and photo-recruitment kinetics.

### Quantifying regularity of SspB-AuroraB clustering on the membrane

To quantify the regularity of SspB-AuroraB patterns on the GUV surface (Fig. 3c,d and Supplementary Fig. 3f,g) we employed generalized recurrence quantification analysis ^30,51,52^. A recurrence plot (RP) is an advanced technique of nonlinear data analysis that represents a two-dimensional binary diagram indicating the recurrence that occur in an *m*-dimensional phase space within a fixed threshold *ε* at different times *i,j*. For one-dimensional time series, the RP is expressed as a square matrix of zeros (non-occurrences) and ones (occurrences) of states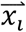 and 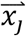 of the system: 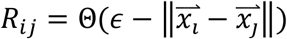, where 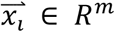, i,j=1,‥N, N is the number of measured states 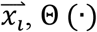 is the step function and ‖·‖ is a norm. To analyze the regularity of SspB-AuroraB on the surface of the GUV, we used an extension of this analysis for spatial data ^56,57^. For a *d*-dimensional Cartesian space, 2 × *d* dimensional spatial recurrence matrix can be defined by: 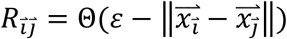, where 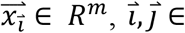 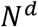 (example projections in Supplementary Fig. 3g). The recurrence quantifications are done based on the diagonal hyper-surfaces defined as surfaces of dimension *N*^*d*^ which are parallel to the *hyper-surface of identity* (defined by, 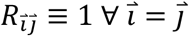). The size of the diagonal hyper-surfaces, 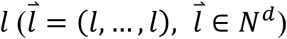 defined by 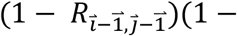 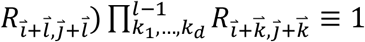, were calculated, and a distribution of surface sizes (*P*(*l*)) was used to calculate the Information entropy. The Information entropy (ENTR) is defined by 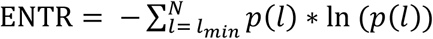, where ln(.) is the natural logarithm, *p*(*l*) is the probability of a diagonal hyper-surface with length *l*, such that 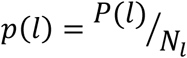 and *N*_*l*_ is the total number of diagonal hyper-surfaces with length greater than a fixed minimum length *l*_*min*_.

The 2D maps of the GUV surface were projected from confocal stacks using the Map3-2D software ^58^. From the spatial map of the SspB-AuroraB on the GUV surface (Fig. 3c, Supplementary Fig. 3f), a rectangular region was selected. After inspecting the corresponding lipid channel, the regions of SspB-AuroraB aggregation were identified by thresholding and substituted with NaNs. Spatial recurrence quantification analysis with *ε* = 0.3 and *l*_*min*_ = 3 was performed on the corresponding rectangular images.

To quantify the SspB-AuroraB fraction in lumen, the 3D z-stack of the GUVs were used. For each stack, the membrane and the lumen were separately masked. The mean intensity per pixel in the 3D lumen (*C*_*lumen*_*mean*_) and on the 2D membrane (*C*_*membrane*_*mean*_) were calculated using imageJ. SspB-AuroraB fraction in the lumen (*C*_*lumen*/*total*_) was calculated using:

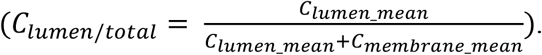

### COPY° characterization

Gly-Stathmin-Cys was sequentially labeled with two organic dyes to generate the sensor COPY° (° = organic). First, the acceptor dye was conjugated to the C-terminal cysteine. Typically, 3 mg of Gly-Stathmin-Cys were incubated with 5 mM TCEP for 1 h at room temperature under an argon atmosphere before buffer exchange via a NAP5 column (GE Healthcare) into degassed 50 mM Hepes pH 7.5, 200 mM NaCl, 2 mM MgCl_2_. A 10-fold molar excess of Atto655-maleimide (Atto-tec) was added and the mixture incubated for 3 h at room temperature under an argon atmosphere. The reaction was stopped by adding 5 mM DTT. Next, without further purification, the donor dye Atto532 (Atto-tec) was conjugated to the N-terminus as described in the section “Protein labeling”. Another sensor version was constructed with Atto647N (Atto-tec) as acceptor dye, which we termed COPY^o2^.

Initial characterization was carried out by evaluating sensitized emission (Supplementary Fig. 4a). Fluorescence emission spectra were measured with a QuantaMaster fluorescence system (PTI) (exc. 520 nm, em. 530 – 720 nm). 1 μM sensor was in 80 mM Pipes pH 6.8, 75 mM KCl, 1 mM EGTA, 2 mM MgCl_2_, 20 mM β-mercaptoethanol. 10 μM tubulin was added to measure the decrease of sensitized emission.

Detailed characterization of the FRET properties was done by analysis of donor fluorescence lifetime. Confocal FLIM experiments were performed on a Leica SP8 confocal microscope (see “Imaging of microtubule asters and morphogenic state transitions in GUVs”) equipped with the FALCON (FAst Lifetime CONtrast) system. Atto532 was excited at 532 nm with a pulse frequency of 40 MHz and emission was collected with three Leica HyD detectors restricted with an Acousto-Optical Beam Splitter (AOBS) to 542 – 563, 568 – 591, and 591 – 620 nm. Data were collected for the donor-only labeled Stathmin, free COPY° and COPY^o2^ (3 μM), and tubulin-bound sensors (+ 20 μM tubulin) in cDICE base buffer (see “Protein encapsulation in GUVs by cDICE”) + 60 μM GTP, 300 mM sucrose, and additional 1 mM trolox. Photons were collected up to approx. 3000 photons per pixel in the sum of all three detectors. Global analysis of FLIM-FRET was implemented in a custom program written in Python according to a described process ^59,60^, yielding τ_D_ (ns) and τ_DA_ (ns) as well as FRET efficiency. Additionally, all pixel values of the output lifetime images were averaged to obtain the absolute lifetimes of the donor for the free and tubulin-bound sensor states (Supplementary Fig. 4b).

Due to the higher signal amplitude obtainable with COPY^o2^, we preferred this variant for stopped-flow and most kinetic measurements. However, since Atto647N (acceptor dye in COPY^o2^) is positively charged and is known to interact with membranes ^61^, we used COPY° for the measurement of stathmin phosphorylation gradients.

### Stopped-flow measurements

Association and dissociation of COPY^o2^ and tubulin were measured in a stopped flow apparatus (Applied Photophysics, Leatherhead, U.K.) with excitation at 535 nm and a 670/30 nm band-pass filter to monitor sensitized emission changes at 25 °C in 80 mM Pipes pH 6.8, 75 mM KCl, 2 mM MgCl_2_, 20 mM β-mercaptoethanol, 60 μM GTP. COPY^o2^ or phosphorylated COPY^o2^ (see “Time-resolved measurements of AuroraB and λ-PPase activity”) was kept constant at 200 nM for the tubulin titration. Dissociation was measured at 200 nM COPY^o2^ and 4 μM tubulin, displacing COPY^o2^ with 60 μM unlabeled stathmin. Data were fit in Origin (OriginLab), and error propagation for derived constants was calculated by Gaussian equation taking the errors of the fit as a starting point.

### Determination of enzymatic kinetic parameters

Stathmin phosphorylation and dephosphorylation kinetics were measured using the fluorescence lifetime change or ratiometric signal of COPY° or COPY^o2^ as indicated in the respective Fig. legends. Lifetime is an absolute, robust and intensity-independent measure of the phosphorylation state and was thereby almost exclusively used for kinetic measurements. Confocal FLIM measurements were performed as described for “Stathmin FRET sensor generation and characterization” with few modifications. Excitation pulse frequency was 20 MHz. Per time-point 1 – 2 million photons were collected in a 256×256 pixels image. During analysis, 4× binning was applied. Measurements were conducted in cDICE-buffer (see “Protein encapsulation in GUVs by cDICE”) + 60 μM GTP, 300 mM sucrose, – EGTA, and + 0.8 mM MnCl_2_ when PPλ was used.

To measure dephosphorylation kinetics, COPY° was pre-phosphorylated (COPY°-P) with 500 nM AuroraB by incubating with 4 mM ATP for 8 h at room temperature and used without further purification. Residual kinase activity was neglected because AuroraB partially loses its activity during the incubation, and also because PPλ deactivates AuroraB in the final sample for the kinetic measurement. Kinetic traces of dephosphorylation were obtained at varying concentrations of COPY°-P (with tubulin present in at least two-fold excess) and 200 nM PPλ. Values for k_cat_ and K_m_ were determined from a fit to the Michaelis-Menten equation. Reported errors are errors of the fit, and the error for k_cat_/K_m_ was derived by Gaussian error propagation.

Since phosphorylation of stathmin under conditions suitable for Michaelis-Menten analysis was too slow to yield sufficiently stable and analyzable signal amplitudes, we estimated k_cat_/K_m_ from monoexponential fits of single-progress curves. 10 μM COPY^o2^ (+ 20 μM tubulin) was phosphorylated by 500 nM or 1 μM SspB-AuroraB, and the data were fit mono-exponentially. The apparent rate constant was converted to a value corresponding to k_cat_/K_m_ by taking the used kinase concentrations into account and averaging the two obtained values. Fit errors were propagated by the Gaussian equation.

Note that the derived kinetic parameters for PPλ and SspB-AuroraB do not consider the presence of multiple phosphorylation sites on stathmin or how many of them need to be phosphorylated to induce tubulin dissociation. The parameters apply for the catalysis of the overall transition between free and tubulin-bound stathmin, which is our direct readout and also the relevant state change for the derivation of the signaling gradient.

### Measurement of stathmin phosphorylation gradients

Spatial COPY° phosphorylation gradients were imaged and quantified by computing the ratio of Atto655/Atto532 fluorescence emission intensities. First, we derive the relationship between the fluorescence FRET ratio to the fraction of tubulin bound stathmin.

The fluorescence intensity detected at the acceptor channel *F*_*A*_ can be separated into three contributions:

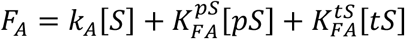

where: *k*_**A**_ is a constant proportional to the fluorophore brightness, [*S*] is the total stathmin concentration, and its product represents the intensity in absence of donor; 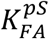 and 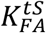 are the contributions to the intensity due to FRET when stathmin is either phosphorylated or tubulin bound, while [*pS*] and [*tS*] are their concentrations, respectively. We assume that stathmin can only adopt these two states and hence [*S*] = [*pS*] + [*tS*].

The fluorescence at the donor channel *F*_*D*_ can be written as:

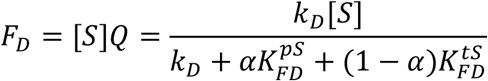

where Q is the quantum yield, *k*_*D*_ is a constant proportional to the donor fluorophore brightness, and 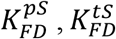 are the contributions to donor quenching by FRET, in the corresponding stathmin states, and *α* = [*pS*]/[*S*] is the fraction of tubulin bound stathmin Normalizing the acceptor fluorescence intensity to the total stathmin concentration yields:

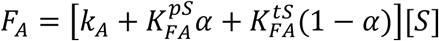

After rearranging terms, we obtain:

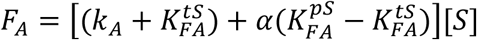

Taking the FRET fluorescence ratio of acceptor fluorescence to donor fluorescence can be then expressed as:

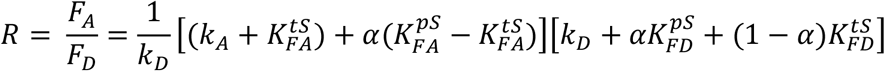

where *R* = *F*_*A*_/*F*_*D*_ is the fluorescence FRET ratio. Rearranging:

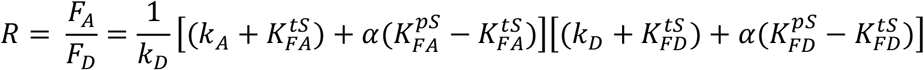

Grouping the remaining constants between parenthesis as,

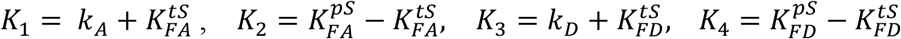

Simplifies the expression to:

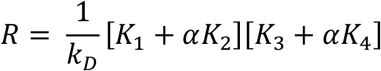

Distributing the product yields:

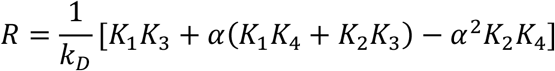

Since FRET is negligible in the tubulin bound state, 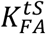 and 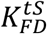 are negligible, and because the FRET efficiency of COPY° is < 30 %, the product 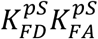 is also negligible since both magnitudes depend on the intrinsic FRET efficiency. Hence, the quadratic term becomes negligible, so we can solve for *α* as:

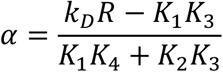

Therefore, the fraction of phospho-stathmin *α* is an approximate linear function of the fluorescence FRET ratio *R*.

The ratiometric intensity is more photon efficient than fluorescence lifetime measurements and therefore provides the sensitivity to detect spatial gradients. Confocal imaging was done on a Leica SP8 microscope (see “Imaging of microtubule asters and morphogenic state transitions in GUVs”). COPY° was excited at 532 nm, and detection was done in photon counting mode, restricted to 542 – 560 nm and 640 – 750 nm for donor and acceptor, respectively. The imaging focus was maintained by Leica Adaptive Focus Control. Imaging conditions were identical when measuring outside of or inside GUVs. The following protein concentrations were used for encapsulation: 40 μM (final) tubulin, 4 μM (final) COPY°, 5 μM C2-iLID, 12 μM SspB-AuroraB (unlabeled), 1 μM λ-PPase. Outside GUVs the following concentrations were used: 20 μM tubulin, 10 μM COPY°, 1 μM iLID_G, 1.5 μM SspB-AuroraB (unlabeled), and varying λ-PPase concentrations as indicated.

For the “dark” state (no iLID activation, no SspB-AuroraB translocation) 40 images at the equatorial plane were taken with 3 s intervals. This spacing was necessary to prevent unwanted iLID activation by 532 nm light. Inside GUVs, this illumination still led to substantial iLID activation, so that no gradient images for the dark state were obtained. Note that Fig. 3a,b also only shows outside gradients in the activated state.

Activation of iLID was done with the 458 nm Argon laser line for the duration of 3 min with 3 s intervals. This sustained activation was performed to allow SspB-AuroraB to undergo full autoactivation and the steady-state of stathmin phosphorylation to be reached. iLID autofluorescence was detected between 568 – 520 nm to be used as a membrane marker during the analysis.

For the activated state, 40 images were taken with minimal intervals. A sequential line scan was performed between iLID activation and COPY° imaging in order to maintain SspB-AuroraB translocation.

In case there was any GUV movement during dark or activated image series acquisition, the frames were aligned in ImageJ using the MultiStackReg plugin based on StackReg ^62^. Then, all images of the series were summed up (more precise than averaging, and made possible by photon counting). We observed an optical xy-aberration between the donor and acceptor channels, which led to asymmetric edge effects around the GUVs in the ratio image. To avoid these, the channels were manually registered to yield a symmetric ratio image calculated as acceptor/donor. Radial profiles were extracted from the ratio images by a custom macro in ImageJ using iLID autofluorescence as a marker to determine the starting position at the membrane and align the profiles of individual GUVs.

Data obtained in this manner were sufficient to describe gradients arising outside of GUVs. Due to lower surface densities of recruited SspB-AuroraB inside GUVs, strong gradients only arose at the positions of SspB-AuroraB clusters. Radial averaging of a single confocal plane thus resulted in profiles with low gradient amplitudes. To visualize the gradient amplitude in cluster vicinity, we measured 7 – 8 μm thick z-stacks of the ratio (1 μm spacing, 10 images per slice) on sufficiently large GUVs at their equatorial plane, so that the GUV diameter would not significantly change throughout the stack. Maximum intensity projections along the z-axis were done to highlight gradient hotspots.

### Paradigmatic model of morphogenesis

The behavior of the two sub-systems, the self-organized morphogenesis of the encapsulated MT-aster as well as the spatial organization of SspB-AuroraB clusters on the GUV membrane, when isolated, can be conceptualized with paradigmatic reaction-diffusion models, where self-amplification of local structures triggers depletion of its free substrate. Generically, this can be represented as:

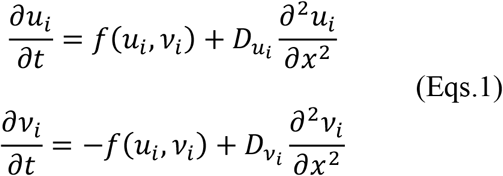

where *u*_*i*_ + *v*_*i*_ = *c*_*i*_ = *const*., *i* = 1,2, and *u*_1_ corresponds to the self-amplified MT-bundles that generate membrane protrusions (Fig. 2e) and *u*_2_ to the SspB-AuroraB clusters on the membrane (Fig. 3e), whereas *v*_1_, represents the astral MTs and *v*_2_-protein monomers. In SynMMS, a joint dynamical system is established, where the protrusion-induced change in the membrane geometry enhances SspB-AuroraB translocation and thereby its activity, that in turn locally promotes the MT-growth (Fig. 5). The joint system is thus described with:

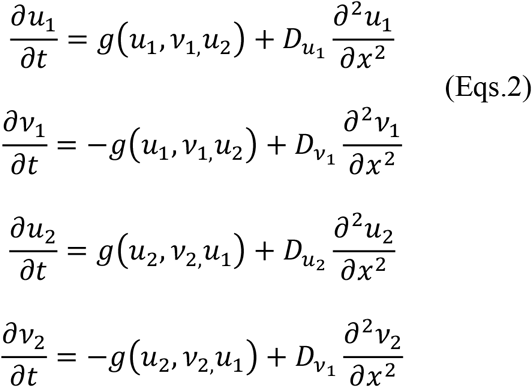

To investigate the dynamical possibilities of this system, we applied Linear Perturbation Analysis (LPA) ^63,64^ that in turn allows to analyze the bifurcation structure of a reaction-diffusion (RD) system. In brief, the method takes advantage of the diffusion discrepancy 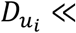 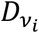 and considering the limit 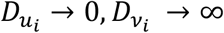, the stability of the homogenous steady state can be probed with respect to a ‘local’ perturbation in the form of a narrow peak of *u*_*i*_ with a negligible total mass, at some location in the domain. In this limit, the localized peak of *u*_*i*_ (*u*L,i) does not influence the surrounding background *u*_*i*_ levels (*u*G,i), whereas due to the ‘infinitely fast’ diffusion, *v*_*i*_ can be described by a single global variable (*v*G,i) that is approximately uniformly distributed in space. This allows writing a set of ODEs that describe the evolution of the perturbation peak as:

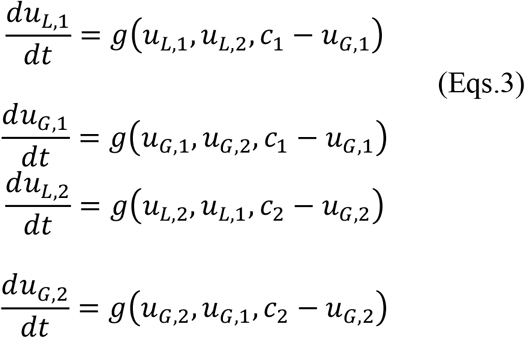

Dynamics of this ODE system (Eqs. 3) under variation of the total SspB-AuroraB concentration (c_2_) was explored using the Xppaut bifurcation software to produce Supplementary Fig. 7a,

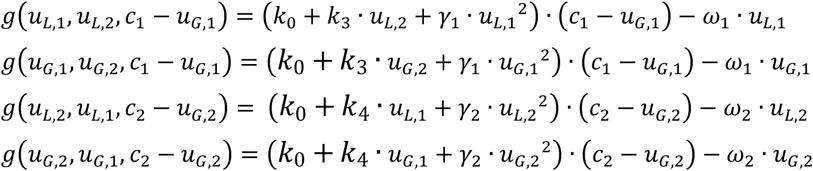

with *k*_0_ = 0.067, *k*_3_ = 1, *k*_4_ = 0.5, *γ*_1_ = 0.4, *γ*_2_ = 5, *ω*_1_ = 0.5, *W*_2_ = 5. The insets in Supplementary Fig. 7a correspond to reaction-diffusions simulations of (Eqs. 2) with different total SspB-AuroraB. From left to right: c_2_=800, 1500, 2750 respectively and c_1_=1000 in all cases.

For the reaction-diffusion simulations in Fig. 5 and Fig. 6, an external signal source corresponding to global light illumination is applied that increases the amount of free AuroraB monomers on the membrane and thereby enhances the cooperative clustering. This is implemented by including ρ which is proportional to the signal strength as ρ · γ_2_ · u_2_^2^. To model the influence of local signal), a Gaussian distribution was used.

For Fig. 5b,g,j and Fig. 6d,h,l all parameters are kept the same, except: for Fig. 5b, c_2_ = (2750, 1500); for Fig. 6l, c_2_ = 2750; for Fig. 6d, c_2_ = 5750, c_1_=2000, *γ*_1_ = 1, *γ*_2_ = 0.5; and for Fig. 5j, c_2_ = 1000, c_1_ = 1500. For Fig. 5b left, c_2_ = 2750 and right, c_2_ = 2750, *γ*_1_ = 1, *γ*_2_ = 0.5. The RD simulations were performed in 1D using a custom-made Python code, with *D*_*u*,*i*_ = 0.01, *D*_*v,i*_ = 40. The integration time was set to 1.5·10^8^, the time step – 7.5·10^−5^, for a total of 10^3^ spatial bins with step size 0.1, and periodic boundary conditions.

### Reaction-diffusion simulation of gradients

We utilize numeric simulations of the reaction diffusion system to visualize the gradients resulting from localizing a kinase to the membrane of a GUV. Our kinetic measurements confirm two tubulin binding sites on stathmin. As the low affinity site (K_D_ = 2 μM for the unphosphorylated, > 200 μM for the phosphorylated state) has a high turnover rate, we consider a simplified setup of five “species”: 1) free tubulin dimers and 2) free stathmin, 3) stathmin tubulin complex (in a 1:1 stoichiometry), 4) phosphorylated stathmin tubulin complex and 5) phosphorylated free stathmin. Here, species 2-5) are actually weakly associated with a second tubulin dimer that slows down effective diffusion, but whose binding state is not strongly influenced by phosphorylation. We assume that the pairwise association/dissociation and kinase/phosphatase reaction can be considered first-order processes governed by a back and a forth rate constant (equivalent to sub-saturation Michaelis-Menten-Kinetics) to reduce the number of parameters in the system. Considering a well-mixed volume (diffusion much faster than reaction), the balance of kinase/phosphatase reaction determines the steady-state levels of phosphorylated stathmin. For our measurements of k_cat_/k_M_ (PPase 0.02 μM^−1^ s^−1^; kinase 0.0011 μM^−1^ s^−1^), at a stoichiometry of 1:1 about 5-10% of stathmin should be phosphorylated with a homogenously flat distribution in the GUV. By recruitment to the membrane, the concentration of the kinase is increased by a factor of up to 500. For this, a 5× increase of fluorescence intensity suffices, as the resolution of ~500 nm dilutes the signal of the “shell” of recruited kinase with a thickness of ~5 nm close to the membrane. Such recruitment leaves a significant fraction of kinase in solution, but due to the unchanged phosphatase activity resulting in a lowered phosphorylated fraction. In a see-saw-like manner, the high kinase activity at the membrane “inverts” phosphorylation after recruitment and leave a fraction of <4% *un*phosphorylated stathmin there. If the phosphatase activity was also recruited, but to the center of the GUV, the result would be a linear gradient completely independent of diffusion, the amplitude of which would solely be determined by the relative strength of kinase and phosphatase activity. In such a case, diffusion would only change the amount of phosphorylated protein transported and how fast the gradient is established. However, as the soluble phosphatase is distributed evenly, the shape of the gradient arises from the interplay of diffusion, phosphatase activity and association/dissociation kinetics of stathmin-tubulin. This defeats any attempt of analytical solutions in more than simplified geometries and requires solving the underlying differential equations numerically.

To this effect, reaction-diffusion simulations are performed by cellular automata approach ^65^ 1D, with a resolution of 0.01 μm per space unit. This corresponds to the kinase homogenously localized to the membrane of a spherical GUV. For a diffusion coefficient of 20 μm^2^ s^−1^ of species 3+4), this translates to a diffusant radius of 3 space units and a time resolution of the simulation of 0.01 s per time step. The diffusant radius of species (1+2+5) is set to 4 space units (corresponding to a slightly too high D = 35 μm^2^ s^−1^, avoiding numerical inconsistencies). Initial conditions were set for all association/dissociation of all species in steady-state without any kinase activity and setting kinase activity to its maximum instantaneously at t_0_. Simulations continue until the relative change of the vectorial sum of all concentrations <10^−9^ results in a steady-state distribution of all species. For dissociation/association reactions, the relevant rate constants are: k_on,stath_ = 16 s^−1^, k_off,stath_ = 0.048 s^−1^, k_on,p-stath_ = 1.8 s^−1^, k_off,p-stath_ = 2.3 s^−1^ and remain fixed across all simulations. In most experimental conditions, 2 – 5 μM of soluble kinase are used with a corresponding activity of k_kin_ = 0.004 s^−1^. We estimate the increase of concentration after translocation to the membrane by a factor of 200 – 500 to yield a kinase activity of k_kin_ = 1 s^−1^ for a depletion of 50% soluble kinase. Conversely, the phosphatase activity at 1 μM can be estimated as k_PPλ_ = 0.02 s^−1^. These parameters produce a spatial distribution of the species (1-5) depicted in Supplementary Fig. 4g (left panel). Higher diffusivity of phospho-stathmin as compared to the tubulin-stathmin-complex (molecular weight 220 kDa, D = 20 μm^2^ s^−1^) leads to a depletion of the total stathmin (free, phosphorylated and tubulin-bound, cyan line in Supplementary Fig. 4g (left panel) at the membrane. This distribution is mirrored by the total tubulin due to its high association rate to stathmin.

Gradients of phosphorylated stathmin and free tubulin result from the competition of kinase activity that effectively “deposits” tubulin near the membrane and entropic redistribution by diffusion as well as rebinding to dephosphorylated unbound stathmin as driven by dephosphorylation. An activity of lambda phosphatase of 0.02 s^−1^ is sufficient to let the gradients decline within 1.5 μm distance from the membrane even at 10000 times faster kinase activity. Interestingly, varying the phosphatase activity for a kinase activity of 1 s^−1^ in agreement with our measurements shows an optimally steep gradient for 100 times slower phosphatase activity. Low phosphatase activity can saturate the GUV with phospho-stathmin and therefore free tubulin due to the shift in partitioning and yields a flat gradient with a high off-set. The highest phosphatase activity abolishes the phospho-stathmin gradient (zero amplitude and offset). Due to the association rate to (phospho-)stathmin being about 20 times faster than even kinase activity, the gradient of free tubulin is “locked” to the phospho-stathmin gradient (Supplementary Fig. 4f).

We additionally perform simulations in 2D to assess the impact of local geometry of protrusions on the gradients of free tubulin. Protrusions feature a much higher surface-to-enclosed-volume ratio compared to undeformed spherical GUVs. Assuming a homogeneous distribution of kinase activity per μm^2^ of surface, we plot the gradient of free tubulin in a GUV with a radius of 32 μm, featuring protrusions of 250 nm diameter with varying lengths (0.5 μm – 22.5 μm). The kinetic parameters were kept identical to previous 1D simulation, specifically with the activities of phosphatase (0.02 s^−1^) and kinase (0.004 s^−1^ homogenously in the medium and 1 s^− 1^ homogenously at membrane).

#### CODE AVAILABILITY

All custom-made code is available upon request to the corresponding author.

