## Supplementary figures and tables for "A synthetic morphogenic membrane system that responds with self-organized shape changes to local light cues"

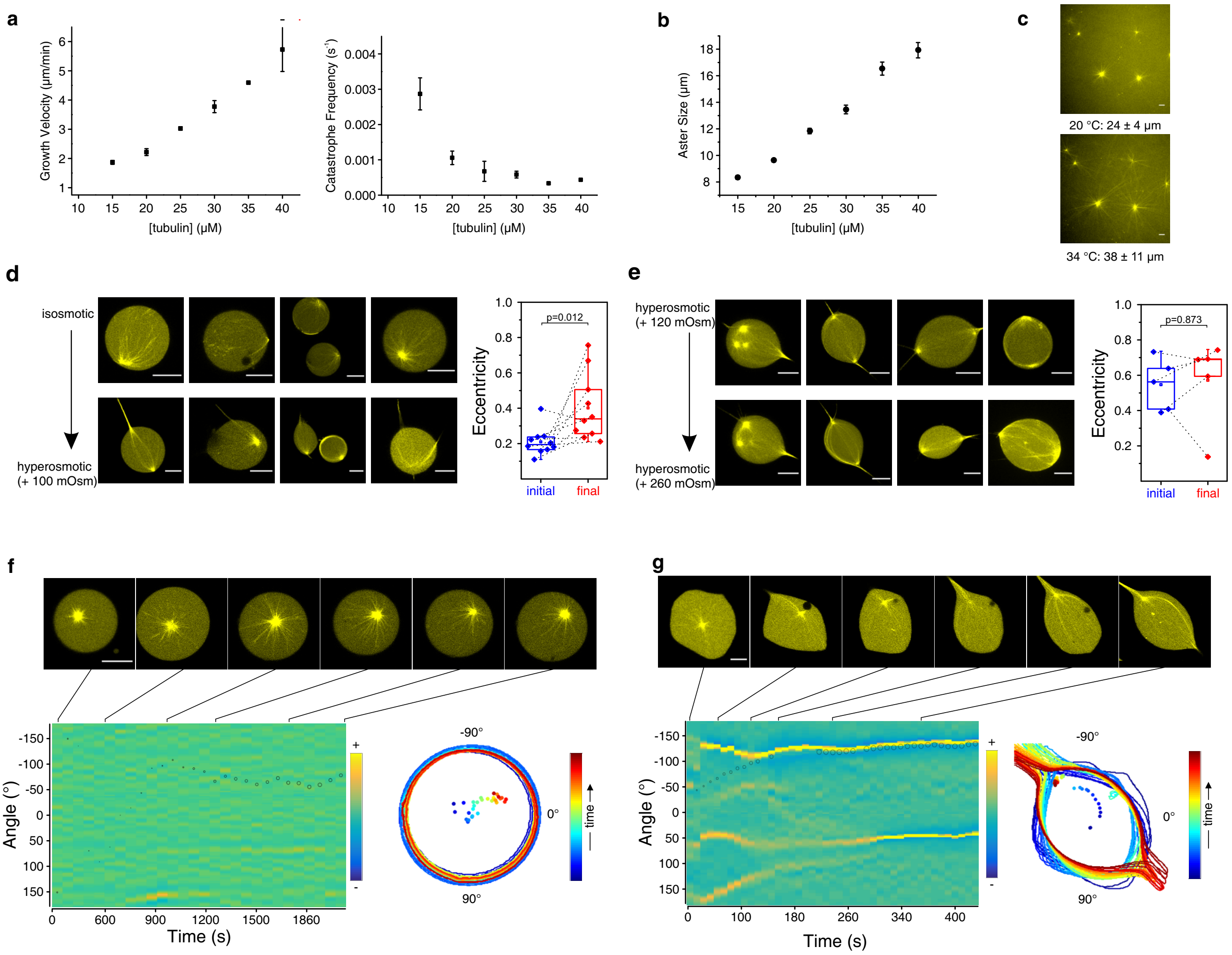


**Supplementary Fig. 1. Characterization of MT dynamics in solution and behavior of MT-asters encapsulated in GUVs.**
(a) Dependence of MT growth velocity (left) and catastrophe frequency (right) on tubulin concentration as determined by single-filament TIRF microscopy assays. Error bars correspond to S.E.M. (> 75 tracked filaments per condition). (b) MT-aster size in bulk as a function of tubulin concentration (>30 asters per condition, error bars are standard error of the fit to the radial intensity profile of the overlaid asters, see Methods). (c) MT-aster size dependency on temperature. Example MT-asters (40 µM tubulin) at 20 °C (top) and at 34 °C (bottom). Average size +/- S.D. is indicated. (d) Left: spherical to polar morphological transitions induced by membrane tension decrease caused by changing outside osmolarity (Methods). Top and bottom rows show corresponding GUVs before and after osmolarity change. Right: corresponding morphometric quantification of the shape by eccentricity at the plane of the centrosome. Dashed lines connect initial and final states of single GUVs. (e) Left: morphological states of initially polar GUVs before (top) and after (bottom) increase in outside osmolarity. Right: corresponding morphometric quantification as in (d). (f) Time-lapse of temperature-induced aster growth in GUVs with a rigid membrane (isosmotic). Top: representative images from the time-lapse, bottom left: kymograph of local membrane curvature with overlaid centrosome position (small circle: central, large circle: peripheral), bottom right: GUV contours during time-lapse, color coded by time progression (dots: corresponding centrosome positions). Lines connect images with the corresponding time in the kymograph. (g) Time-lapse of temperature-induced aster growth in a GUV with low membrane tension. Representation as in (f). p-values from two-sample Kolmogorov-Smirnov test. Scale bars are 10 µm.

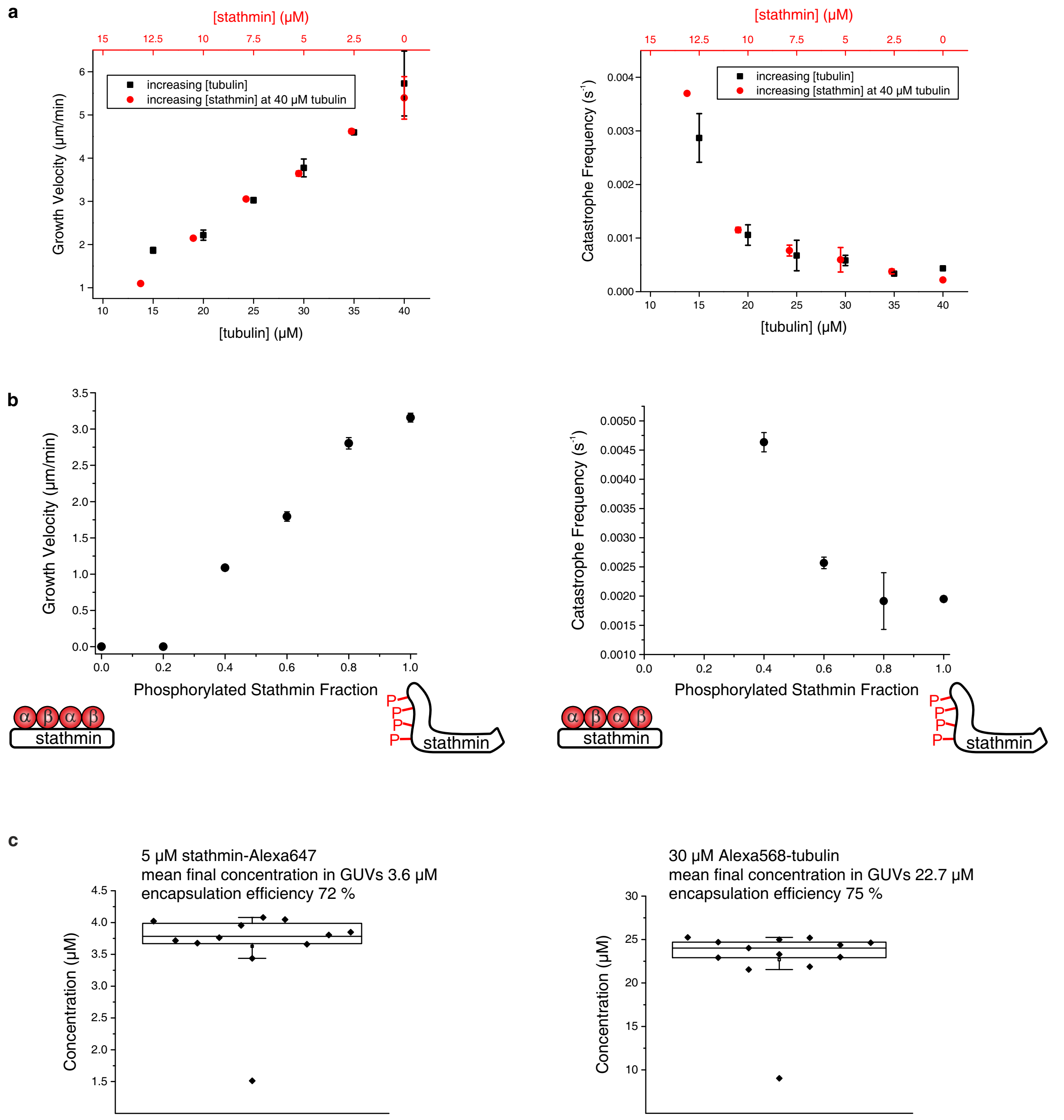


**Supplementary Fig. 2. Stathmin regulation of MT-dynamics.**

(a) Dependence of MT growth velocity and catastrophe frequency on tubulin and stathmin concentration as determined by single-filament TIRF microscopy assays, measured at 40 µM tubulin with increasing stathmin concentrations (red axis on top). Data for tubulin in the absence of stathmin (black) is identical to those in Supplementary Fig. 1a. Introducing stathmin into the system is equivalent to decreasing tubulin concentration, assuming a 1:2 binding stoichiometry (stathmin:tubulin). Error bars correspond to S.E.M. (> 75 tracked filaments per condition). (b) Dependence of MT growth velocity (left) and catastrophe frequency (right) at 40 µM tubulin on the fraction of pStathmin over total stathmin determined by single-filament TIRF microscopy assays. The fraction of pStathmin was varied keeping a fixed total concentration of 20 µM Stathmin. Error bars correspond to S.E.M. (> 75 tracked filaments per condition). (c) Quantification of encapsulation efficiency of stathmin-Alexa647 (labeled via maleimide as described for FRET sensor generation in Methods) and Alexa568-tubulin. Left: box plot of encapsulated stathmin-Alexa647 concentrations from 5 µM stock solution. The mean value was 3.6 ± 0.7 µM, corresponding to 72 ± 14 % encapsulation efficiency. For quantification, 1 µM stathmin-Alexa647 was added to the outside as reference. Right: box plot of encapsulated Alexa568-tubulin concentrations from 30 µM stock solution. The mean value was 22.7 ± 4 µM, corresponding to 75 ± 13 % encapsulation efficiency. For quantification, 1 µM Alexa568-tubulin was added to the outside as reference.

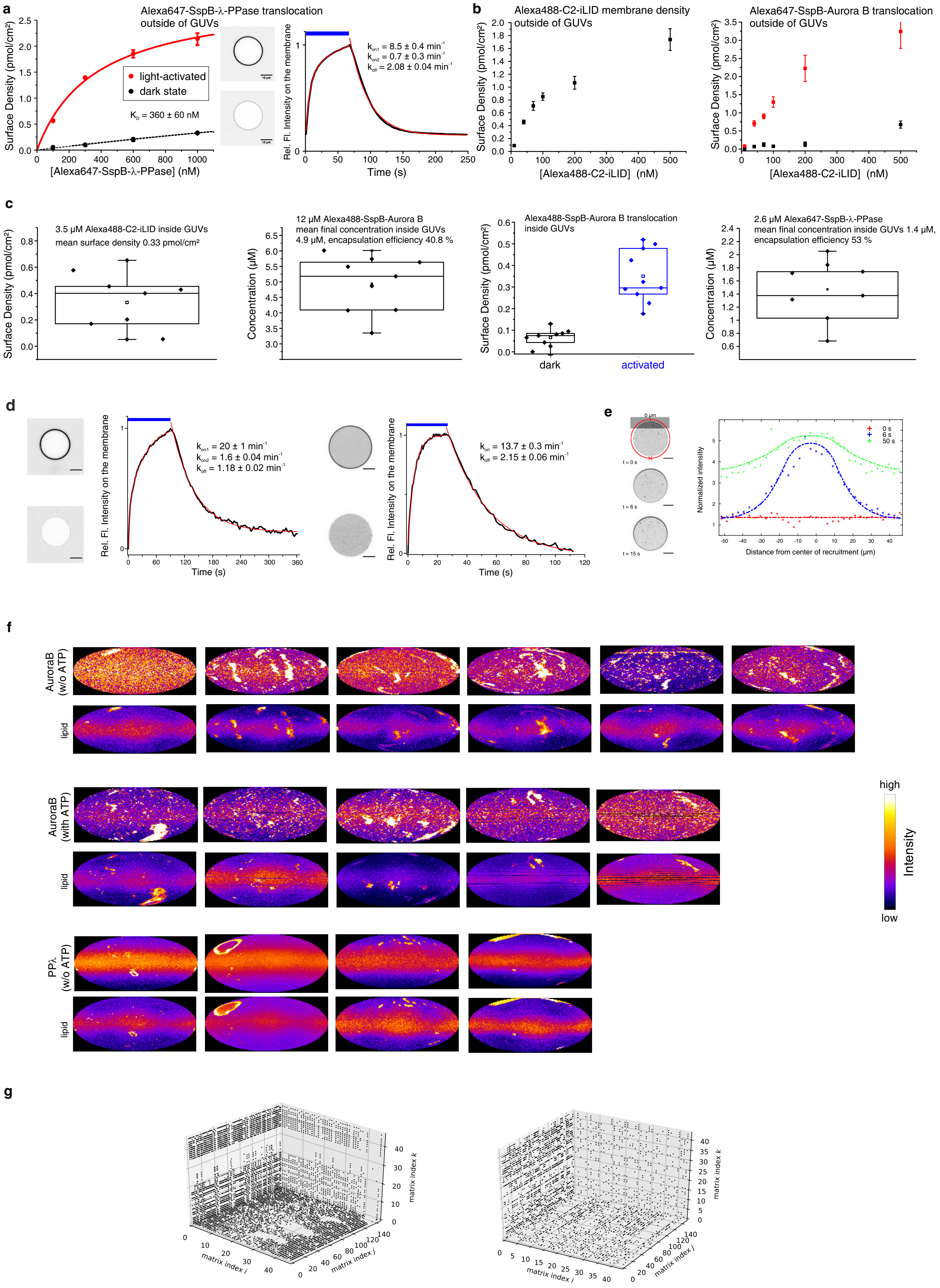
a
Alexa647-SspB-λ-PPase translocation outside of GUVs
light-activated
dark state
K_D = 360 ± 60 nM
Surface Density (pmol/cm²)
[Alexa647-SspB-λ-PPase] (nM)
Rel. Fl. Intensity on the membrane
k_on1 = 8.5 ± 0.4 min⁻¹
k_on2 = 0.7 ± 0.3 min⁻¹
k_off = 2.08 ± 0.04 min⁻¹
Time (s)
b
Alexa488-C2-iLID membrane density outside of GUVs
[Alexa488-C2-iLID] (nM)
Alexa647-SspB-Aurora B translocation outside of GUVs
c
3.5 µM Alexa488-C2-iLID inside GUVs
mean surface density 0.33 pmol/cm²
12 µM Alexa488-SspB-Aurora B
mean final concentration inside GUVs 4.9 µM, encapsulation efficiency 40.8 %
Concentration (µM)
Alexa488-SspB-Aurora B translocation inside GUVs
dark
activated
2.6 µM Alexa647-SspB-λ-PPase
mean final concentration inside GUVs 1.4 µM, encapsulation efficiency 53 %
d
k_on1 = 20 ± 1 min⁻¹
k_on2 = 1.6 ± 0.04 min⁻¹
k_off = 1.18 ± 0.02 min⁻¹
k_on = 13.7 ± 0.3 min⁻¹
k_off = 2.15 ± 0.06 min⁻¹
e
0 µm
t = 0 s
t = 6 s
t = 15 s
0 s
6 s
50 s
Normalized intensity
Distance from center of recruitment (µm)
f
AuroraB (w/o ATP)
lipid
AuroraB (with ATP)
PPλ (w/o ATP)
high
Intensity
low
g
matrix index i
matrix index j
matrix index k

**Supplementary Fig. 3. Light-mediated SspB translocation and cDICE encapsulation efficiency.**

(a) Left: Alexa647-SspB-PPλ surface density at the membrane (Methods) on the outside of GUVs. The construct increases its affinity for membrane-tethered Alexa488-C2-iLID upon blue light illumination and thereby translocates to the membrane. The surface density in the light-activated state (red) was fit to a binding isotherm, yielding a $K_D = 360 \pm 60$ nM. A surface density of about 2 pmol/cm² could be achieved with low dark binding (black). Datapoints are mean ± S.D. of 3 GUVs. Center: example images of Alexa647-SspB-PPλ at 600 nM (excitation: 650 nm, detection 705 ± 45 nm) without photo-activation (bottom) and activated by blue light (488 nm) (top). Right: kinetics of blue light-induced membrane recruitment and release after switching off the photo-activation light. The data were fitted by bi- and monoexponential functions, respectively, yielding apparent rate constants $k_{recruit1} = 8.5 \pm 0.4$ min$^{-1}$, $k_{recruit2} = 0.7 \pm 0.3$ min$^{-1}$, $k_{release} = 2.08 \pm 0.04$ min$^{-1}$. The kinetic data is an average from 6 GUVs. Kinetics were acquired with 600 nM Alexa647-SspB-PPλ and 100 nM C2-iLID. Blue bar: blue light irradiation period. (b) Left: titration of Alexa488-C2-iLID to outside GUV membranes yielding surface densities of up to 2 pmol/cm². Right: light-induced recruitment (red) of Alexa647-SspB-AuroraB (1.5 µM in bulk) for different concentrations of Alexa488-C2-iLID. Alexa647-SspB-AuroraB binding without blue photo-activation light (black). Datapoints depict mean ± S.D. of 3 GUVs. (c) Quantification of encapsulation efficiency and membrane densities of Alexa488-C2-iLID, Alexa647-SspB-AuroraB, and Alexa647-SspB-PPλ (from left to right). First plot: box plot of Alexa488-C2-iLID surface density upon encapsulation of 3.5 µM protein. The mean value ± S.D. was 0.33 ± 0.2 pmol/cm², which is 3 – 6-fold lower than outside GUVs. 1 µM soluble Alexa488-iLID-tRac1 was added to the outside for quantification, and the brightness ratio of the two proteins was determined separately. Second plot: box plot of Alexa488-SspB-AuroraB concentrations upon encapsulation of 12 µM protein. The mean ± S.D. value was 4.9 ± 0.9 µM, corresponding to 40.8 ± 7.5 % encapsulation efficiency. For quantification, 300 nM Alexa488-SspB-AuroraB was added to the outside as reference. Third plot: box plots of Alexa488-SspB-AuroraB surface density inside GUVs before and after photo-activation. 0.35 ± 0.12 pmol/cm² (mean ± S.D.) were on average recruited upon activation. Fourth plot: box plot of Alexa647-SspB-PPλ concentrations upon encapsulation of 2.6 µM protein. The mean value was 1.40 ± 0.45 µM, corresponding to 53 ± 17 % encapsulation efficiency. (d) Kinetics of blue light-induced Alexa647-SspB-AuroraB membrane recruitment and release after switching off photo-activating light, on the outside (left) and inside (right, same graph as in Fig. 3b) of GUVs. The obtained apparent rate constants are for outside: $k_{recruit1} = 20 \pm 1$ min$^{-1}$, $k_{recruit2} = 1.60 \pm 0.04$ min$^{-1}$, $k_{release} = 1.18 \pm 0.02$ min$^{-1}$, and for inside: $k_{recruit} = 13.7 \pm 0.3$ min$^{-1}$, $k_{release} = 2.15 \pm 0.06$ min$^{-1}$. The protein concentrations outside were 1.5 µM Alexa647-SspB-AuroraB and 1 µM iLID_G. Protein concentrations for encapsulation were 2 µM Alexa647-SspB-AuroraB and 80 µM iLID_G. The average of 4-6 GUVs for each condition is shown. Example images of GUVs pre- and post-translocation were taken at 1.5 µM Alexa647-SspB-AuroraB, 200 nM Alexa488-C2-iLID outside GUVs and 2 µM Alexa647-SspB-AuroraB, 40 µM iLID_G inside GUVs. (e) Local photo-recruitment of Alexa647-SspB-PPλ to C2-LID. The ROI (gray shaded area) was irradiated with blue light for local photo-recruitment for 50 s. Left: snapshots of the time-lapse of Alexa647-SspB-PPλ before (0 s) and during (6 s, 15 s) activation. Red arrow indicates distance along the membrane from the center of photo-recruitment. Right: resulting Alexa647-SspB-PPλ intensity distribution along the membrane before (0 s, red), during (6 s, blue) and at the end (50 s, green) of activation. Profiles were fitted by 2D CA-simulation, which revealed a coefficient of effective lateral diffusion of 26.4 µm²/s on the membrane. +: fluorescence intensity, line: simulation fit. (f) Surface intensity maps of GUVs with membrane-translocated SspB-AuroraB in absence (top panel) or presence (middle panel) of ATP and membrane-translocated SspB-PPλ in absence of ATP (bottom panel). For each case, lower row shows the corresponding fluorescent LissamineRhodamineB-DOPE lipid distribution. The horizontal stripes in the surface maps are result of the projection, and occur when the GUV is large as compared to confocal z-spacing. (g) Examples of spatial recurrence plots (RP) for the intensity distribution of membrane-translocated Alexa488-SspB-AuroraB (left) or Alexa647-SspB-PPλ (right), used for the quantification in Fig. 3d. Shown are three-dimensional slices of the four-dimensional RPs. Scale bars are 10 µm

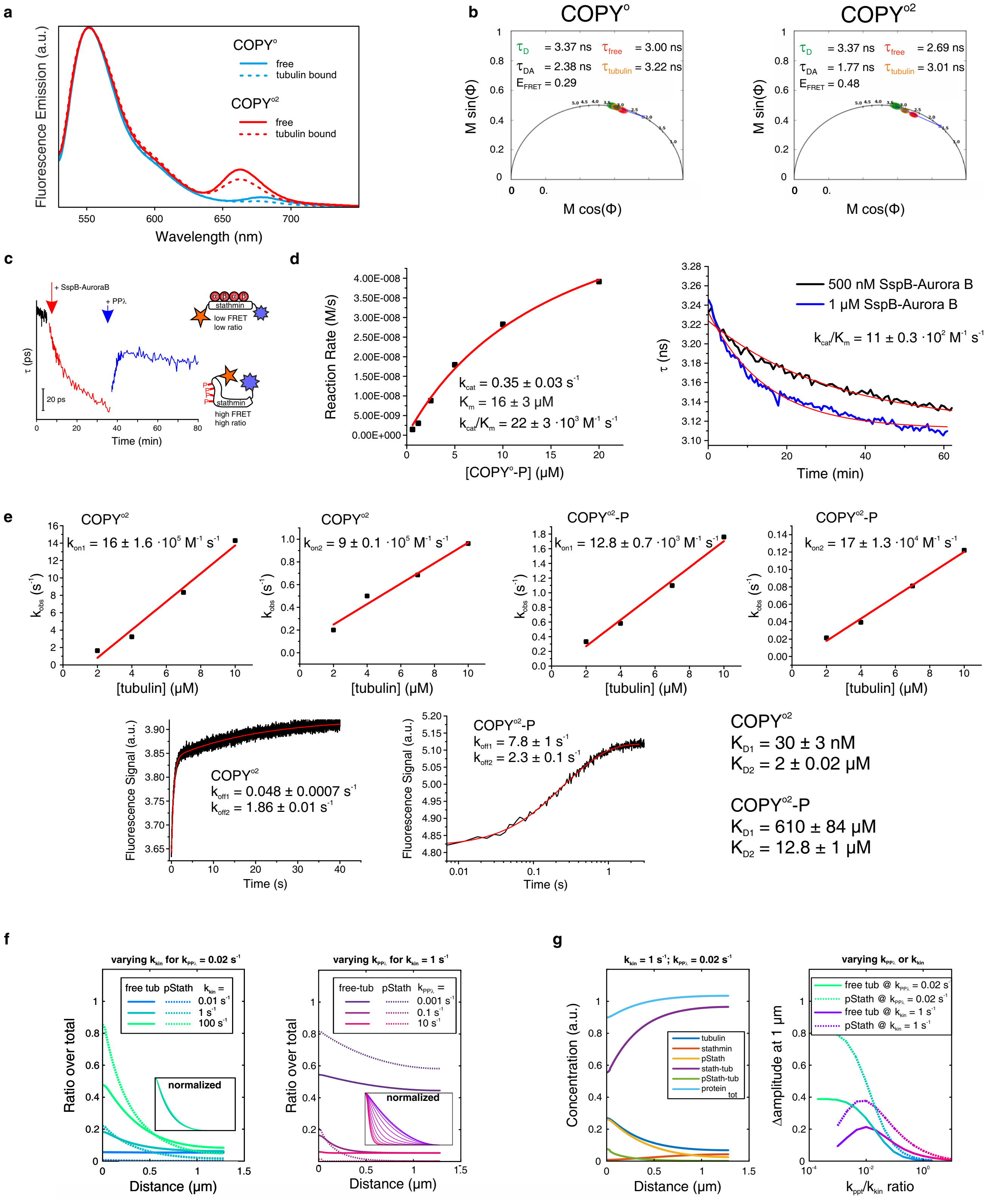


**Supplementary Fig. 4. Photophysical characterization of the stathmin conformational FRET sensor COPY°, kinetics of its interaction with tubulin and simulations of tubulin gradient.**

(a) Fluorescence emission spectra of COPY° and COPY°2 in absence (solid lines) or presence tubulin (dashed lines). COPY° and COPY°2 both have Atto532 as donor and Atto655 or Atto647N as acceptor, respectively. Atto532 was excited at 520 nm. Acceptor sensitized emission peaks are visible between 650 and 700 nm. (b) Quantification of COPY° and COPY°2 FRET efficiency by Time-Correlated Single Photon Counting Fluorescence Lifetime Imaging Microscopy (TCSPC-FLIM). Phasor plot representation of the Fourier components of the first harmonic frequency (20 MHz) of TCSPC-FLIM data is shown. Fluorescence lifetime of the $COPY^{o}$ and $COPY^{o2}$ constructs was determined by global analysis of FLIM data. The fluorescence lifetime of $COPY^{o}$ and $COPY^{o2}$ without acceptor ($\tau_{D}$) was 3.37 ns (green). $COPY^{o}$ and $COPY^{o2}$ exhibited lifetimes with corresponding FRET efficiencies ($\tau_{DA}$ ± S.D. of the lifetime image, $E_{FRET}$ ± propagated error) of 3.0 ± 0.1 ns ($E_{FRET}$: 10.98 ± 0.03 %) and 2.69 ± 0.08 ns ($E_{FRET}$: 20.18 ± 0.02 %) in the absence of tubulin (red), while tubulin binding (orange) increased the lifetimes to 3.2 ± 0.1 ($E_{FRET}$: 5.04 ± 0.03 %) and 3.01 ± 0.09 ns ($E_{FRET}$: 10.68 ± 0.02 %), respectively. (c) Fluorescence lifetime measurement of $COPY^{o}$ phosphorylation and dephosphorylation in bulk (10 µM $COPY^{o}$, 20 µM tubulin, 2 µM SspB-AuroraB, 500 nM PPλ), which is analogous to the ratiometric measurement in Fig. 3f. Enzyme addition indicated by arrows. Right: scheme of the FRET-sensor $COPY^{o}$ as in Fig. 3f. (d) Kinetic parameters of $COPY^{o2}$ dephosphorylation by PPλ and phosphorylation by SspB-AuroraB. Left: Michaelis-Menten analysis of $COPY^{o}$ ($COPY^{o}$-P, Methods, Supplementary Table 1) dephosphorylation by PPλ. Right: $COPY^{o2}$ (10 µM) phosphorylation kinetics by SspB-AuroraB in the presence of 20 µM tubulin. $k_{cat}/K_{m}$ was estimated from corresponding monoexponential fits at two SspB-AuroraB concentrations (Supplementary Table 1). (e) Kinetic and binding parameters of the $COPY^{o2}$-tubulin interaction as measured by sensitized emission in a stopped flow apparatus. Top row: linear fits of apparent association rate constants of $COPY^{o2}$ and $COPY^{o2}$-P to tubulin. Two association rate constants corresponding to two binding sites were determined. Bottom row: biexponential fits of tubulin dissociation from $COPY^{o2}$ and $COPY^{o2}$-P, obtaining two dissociation rate constants. From these rate constants, corresponding affinities were calculated (Supplementary Table 1). (f) 1D reaction-diffusion simulations of pStathmin and tubulin gradients. Stathmin association/dissociation parameters were set to the values measured above (Methods). $k_{PP\lambda}$: dephosphorylation reaction rate, $k_{kin}$: phosphorylation reaction rate. Left: free tubulin (solid lines) and the fraction of free phosphorylated stathmin (dashed lines) are plotted against distance to the membrane for $k_{PP\lambda} = 0.02\ s^{-1}$ and $k_{kin}$ varying between 0.01 and 100 $s^{-1}$ ($k_{kin} = 1\ s^{-1}$). The inset represents the normalized free tubulin gradients. Right: free tubulin (solid lines) and the fraction of free phosphorylated stathmin (dashed lines) are plotted against distance to the membrane for $k_{kin} = 1\ s^{-1}$ and $k_{PP\lambda}$ varying between 0.001 and 10 $s^{-1}$. The inset represents the normalized free tubulin gradients, showing that the decay length changes with changing phosphatase reaction rate, but is robust against kinase reaction rate. (g) Left: Concentration spatial profiles of the five species (box legend) in steady-state, shown here up to a distance of 1.3 µm in a 30 µm diameter GUV. Right: Difference between maximum amplitude at the membrane, 0 µm, and at 1 µm distance (Δamplitude). The amplitude difference is maximal when the kinase activity is 100 times larger than the phosphatase activity.

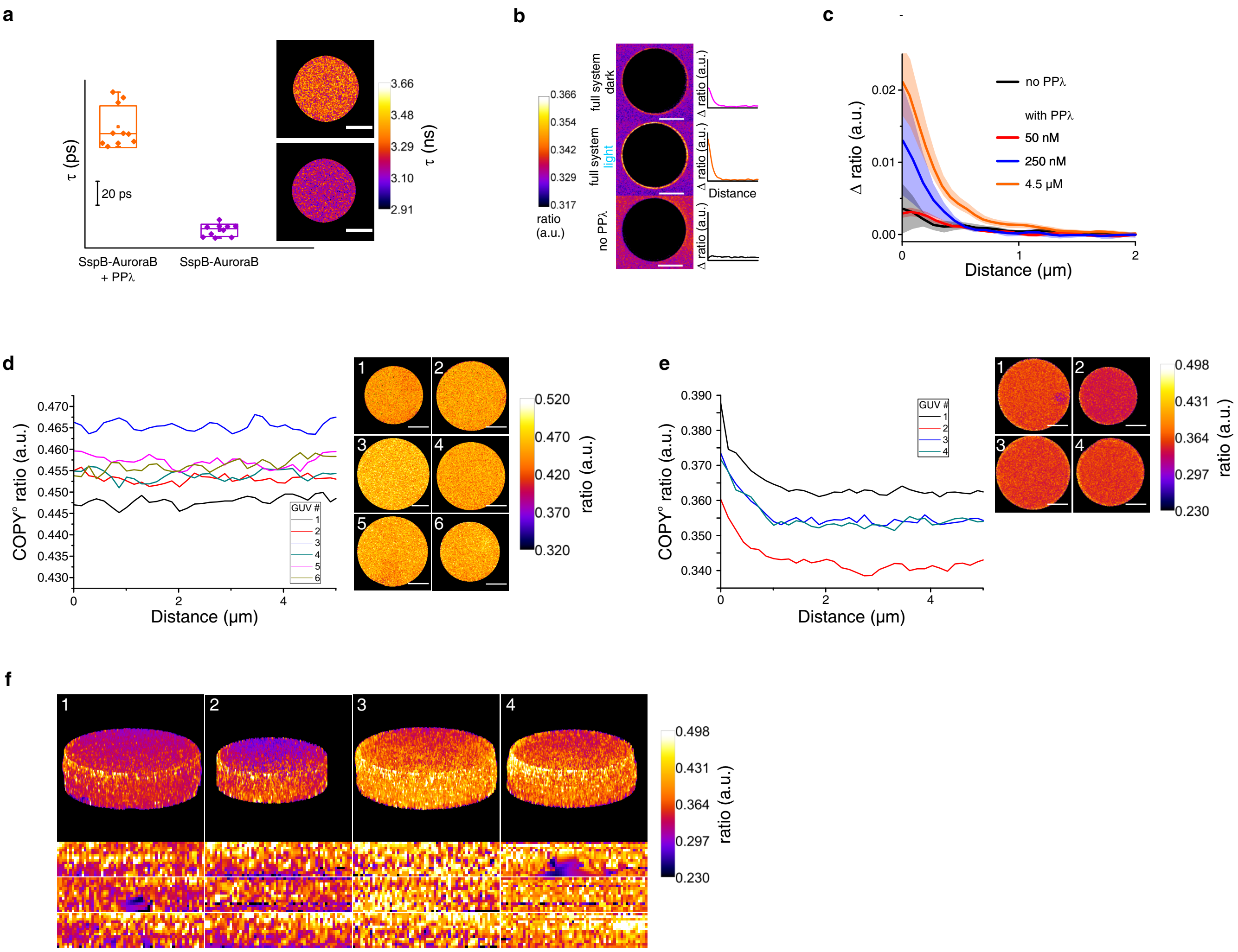


**Supplementary Fig. 5. Stathmin phosphorylation gradient.**
(a) COPY° fluorescence lifetimes (τ) in GUVs with PPλ (orange, Methods) or in the absence of PPλ (purple). Example images to the right. In the absence of PPλ a lower lifetime indicates a higher proportion of pStathmin. (b) Ratiometric images of COPY° outside of GUVs with iLID on the membrane, SspB-AuroraB and tubulin in the bulk (concentrations in Methods). Top: +250 nM PPλ in the absence of blue photo-activating light, center: after light-induced translocation of SspB-AuroraB, bottom: without PPλ. Corresponding bulk ratio subtracted radial profiles (Δratio) to the right. (c) Quantification of Δratio gradients with different PPλ concentrations (mean ± S.E.M. of 4 GUVs for each condition). (d) Plots of COPY° FRET ratio profiles (0 μm defines the membrane position) and corresponding images derived from maximum intensity ratio z-stacks (8 slices at equatorial plane) inside GUVs without PPλ (Methods). Concentration of components were: 40 μM (final) tubulin, 4 μM (final) COPY°, 5 μM C2-iLID and 12 μM SspB-AuroraB. (e) Plots of COPY° FRET ratio profiles and corresponding images for full encapsulated signaling module: 1 μM PPλ, other concentrations as in (d). Note that the steady-state ratio in the GUVs beyond 2 μm (lumen) is lower than without PPλ (d), indicating that PPλ dominates over the AuroraB kinase activity in the lumen before SspB-AuroraB translocation, maintaining stathmin in a dephosphorylated steady state. (f) Top: 3D projections of the GUVs corresponding to the ratio profiles in (e), and bottom: projected 2D surface maps of the ratio in membrane vicinity. Scale bars are 10 μm.

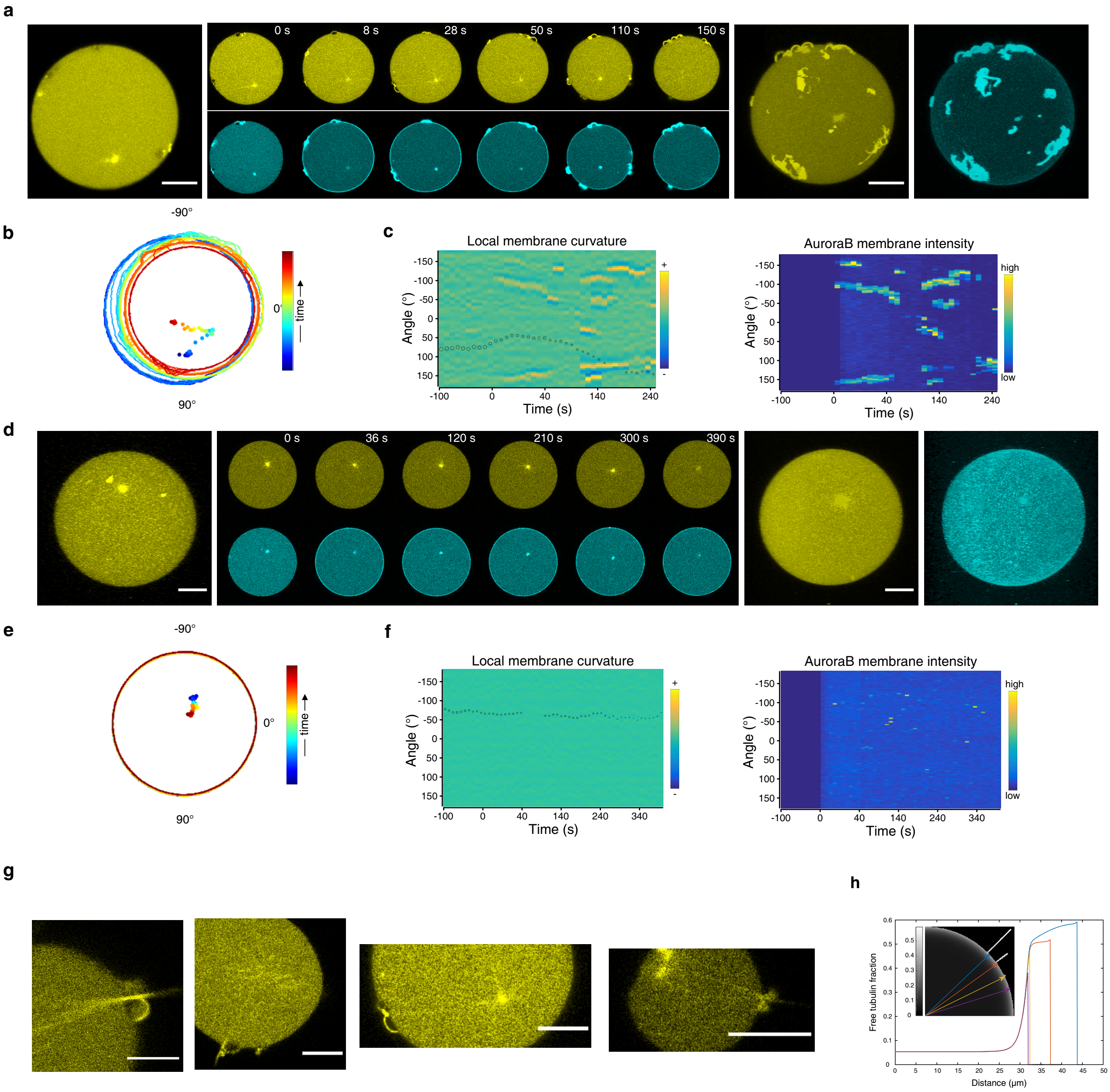


**Supplementary Fig. 6. Feedback between astral MT growth and signaling at the deformable membrane.**

(a) Time lapse of Alexa647-tubulin (yellow) and Alexa488-SspB-AuroraB (cyan) at the equatorial plane of a SynMMS with an initially small MT-aster during continuous global illumination with blue light. Maximum intensity projections of Alexa647-tubulin fluorescence of a confocal z-stack obtained before (left) and after the time lapse series (right, together with Alexa488-SspB-AuroraB) are shown. (b) Corresponding GUV contours for the entire time-lapse, color coded by time progression (dots: corresponding centrosome positions). (c) Kymographs of local membrane curvature (left; overlaid with centrosome position, small circle: central, large circle: peripheral) and corresponding Alexa488-SspB-AuroraB membrane intensity (right) for SynMMS in (a). Negative time: prior to activation. (d) Time lapse as in (a) for a control SynMMS$^{-stat}$ (without encapsulated stathmin) with an initially small MT-aster during continuous global illumination with blue light. Corresponding contours (e) and kymographs (f) for (d). (g) Alexa647-tubulin confocal fluorescence micrographs of SynMMS showing the connection of astral MTs to branching MT-protrusions. (h) Free tubulin gradient generated by stathmin phosphorylation cycle illustrated by 2D simulation of a GUV with thin protrusions (Methods). The enhanced gradient amplitude in protrusions further increases with their length. Colored 1D-profiles taken along lines as indicated by corresponding arrows in the inset.

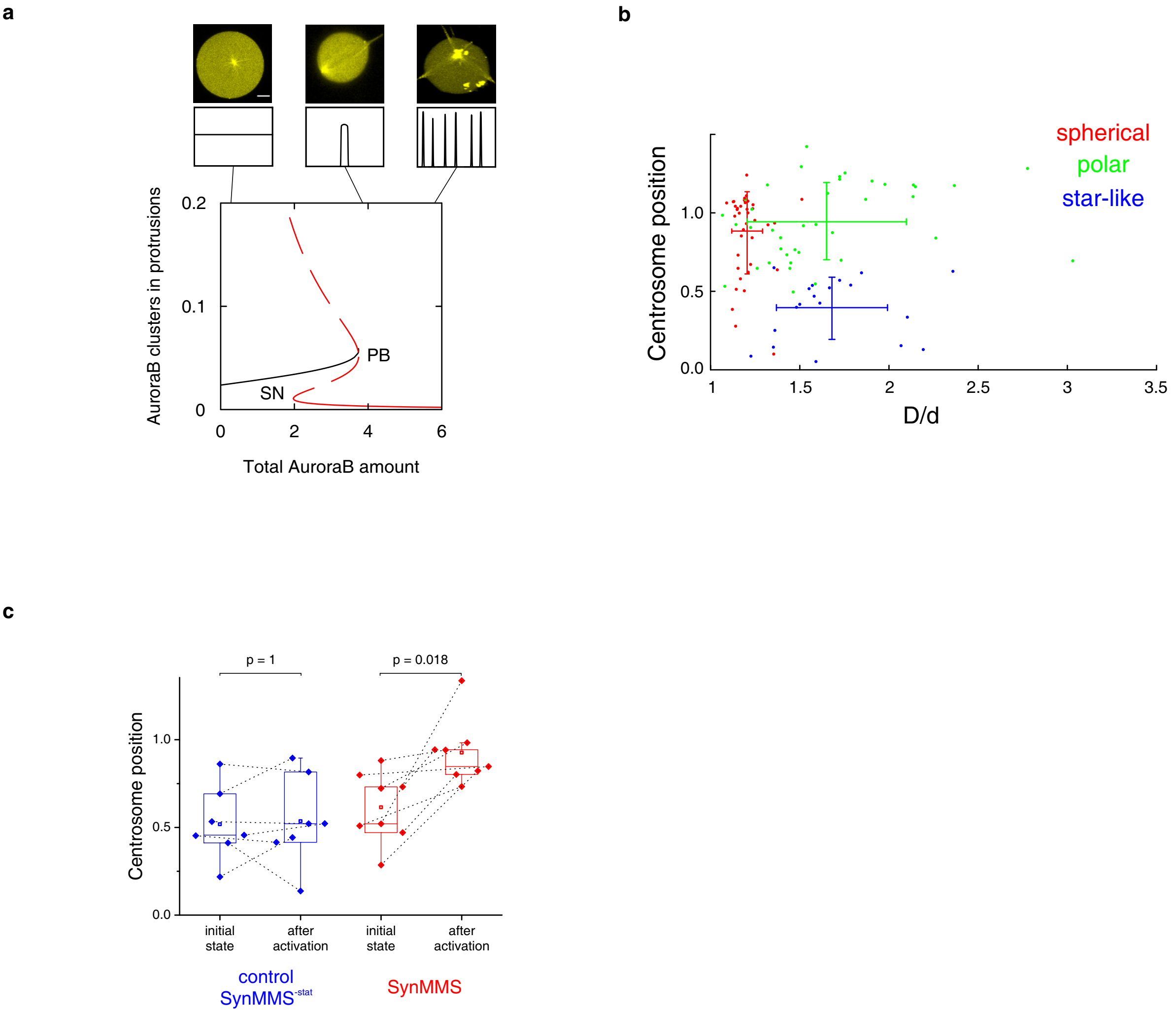


**Supplementary Fig. 7. Dynamical features of SynMMS lead to self-organized morphologies depending on signaling activity at the membrane.**

(a) Bifurcation diagram of the generalized substrate-depletion model of the full system depicted in Fig. 5a. An SspB-AuroraB concentration at the membrane lower than that of the symmetry-breaking pitchfork bifurcation leads to a spherical morphology with small aster size, whereas for concentrations above this point, the system either self-organizes to polar (organization in the vicinity of the bifurcation) or to star-like morphologies (after the bifurcation). Bifurcations: PB - pitchfork, SN - saddle-node. Solid lines: stable homogenous (black) and symmetry broken (red) solution. Dashed line: unstable solution. Top: representative images and numerical distribution of spatial structures at indicated parameter positions in the diagram. (b) Morphometric analysis of initial morphologies of SynMMS or encapsulated microtubule-asters. Plot shows centrosome position (centrosome position: 0 – centered, 1 – membrane proximal) versus irregular parameter (D/d: ratio between the diameter of the minimal bounding sphere and the diameter of the maximal inscribed sphere: 1 for perfect spheres and >1 for deformed or protruded shapes) for each GUV (Methods). From these, three morphological classes that occupy different regions in morphometric space could be identified: star-like (blue), polar (green) and spherical (red). The bars represent the S.D. from the average position of each class. (c) Morphometric quantification of centrosome position before (left) and after (right) blue light-induced translocation of SspB-AuroraB for SynMMS (red) and control SynMMS$^{\mathbf{-stat}}$ (blue). In SynMMS, activation leads to MT-growth and thereby centrosome decentering, which does not occur in SynMMS$^{\mathbf{-stat}}$ because of the absence of stathmin. p-values from two sample Kolmogorov-Smirnov test. Scale bars are 10 μm.

**Supplementary Table 1. Kinetic parameters of stathmin phosphorylation/dephosphorylation and interaction with tubulin.**

| | **Stathmin** | **Phospho-Stathmin** |
|---|---|---|
| **Phosphorylation** | | |
| $k_{cat}/K_m$ | $11 \pm 0.3 \cdot 10^2$ $M^{-1}$ $s^{-1}$ | |
| **Dephosphorylation** | | |
| $k_{cat}$ | | $0.35 \pm 0.03$ $s^{-1}$ |
| $K_m$ | | $16 \pm 3$ µM |
| $k_{cat}/K_m$ | | $22 \pm 5 \cdot 10^3$ $M^{-1}$ $s^{-1}$ |
| **Tubulin interaction** | | |
| association rate constants | $k_{on1} = 16 \pm 1.6 \cdot 10^5$ $M^{-1}$ $s^{-1}$<br>$k_{on2} = 9.0 \pm 0.1 \cdot 10^5$ $M^{-1}$ $s^{-1}$ | $k_{on1} = 12.8 \pm 0.7 \cdot 10^3$ $M^{-1}$ $s^{-1}$<br>$k_{on2} = 17 \pm 1.3 \cdot 10^4$ $M^{-1}$ $s^{-1}$ |
| dissociation rate constants | $k_{off1} = 0.0480 \pm 0.0007$ $s^{-1}$<br>$k_{off2} = 1.86 \pm 0.01$ $s^{-1}$ | $k_{off1} = 8 \pm 1$ $s^{-1}$<br>$k_{off2} = 2.3 \pm 0.1$ $s^{-1}$ |
| affinities | $K_{D1} = 30 \pm 3$ nM<br>$K_{D2} = 2.00 \pm 0.02$ µM | $K_{D1} = 610 \pm 84$ µM<br>$K_{D2} = 13 \pm 1$ µM |

### Supplementary Table 2. Amino acid sequences of used recombinant protein constructs.

| Construct details | amino acid sequence |
|---|---|
| **Gly-Stathmin-Cys**<br>pentaglycine, mouse Stathmin Δ1, additional C-terminal cysteine | GGGGGASSDIQVKELEKRASGQAFELILSPRSKESVPDFPLSPPKKKDLSLEEIQKKLEAAEERRKSHEAEV<br>LKQLAEKREHEKEVLQKAIEENNNFSKMAEEKLTHKMEANKENREAQMAAKLERLREKDKHVEEVRKNKES<br>KDPADETEADC |
| **Gly-iLID-tRac1**<br>pentaglycine, SGS linker, iLID, $(GS)_4$ linker, C-terminus of human Rac1 (175-192) | GGGGGSGSLATTLERIEKNFVITDPRLPDNPIIFASDSFLQLTEYSREEILGRNCRFLQGPETDRATVRKIRDA<br>IDNQTEVTVQLINYTKSGKKFWNVFHLQPMRDYKGDVQYFIGVQLDGTERLHGAAEREAVCLIKKTAFQIAE<br>AANDENYFGSGSGSCPPPVKKRKRKCLLL |
| **Gly-C2-iLID**<br>tetraglycine, $(GS)_4$ linker, C2 domain from bovine lactadherin (270-427), $(GS)_3$, helical $(EAAAK)_3$ linker, $(GS)_3$, iLID | GGGGGSGSGSGSCTEPLGLKDNTIPNKQITASSYYKTWGLSAFSWFPYYARLDNQGKFNAWTAQTNSASE<br>WLQIDLGSQKRVTGIITQGARDFGHIQYVAAYRVAYGDDGVTWTEYKDPGASESKIFPGNMDNNSHKKNIF<br>ETPFQARFVRIQPVAWHNRITLRVELLGCGSGSGSEAAAKEAAAKEAAAKGSGSGSLATTLERIEKNFVITD<br>PRLPDNPIIFASDSFLQLTEYSREEILGRNCRFLQGPETDRATVRKIRDAIDNQTEVTVQLINYTKSGKKFWNV<br>FHLQPMRDYKGDVQYFIGVQLDGTERLHGAAEREAVCLIKKTAFQIAEAANDENYF |
| **λ-phosphatase**<br>PPase ORF221 of bacteriophage lambda | GMRYYEKIDGSKYRNIWVVGDLHGCYTNLMNKLDTIGFDNKKDLLISVGDLVDRGAENVECLELITFPWFRA<br>VRGNHEQMMIDGLSERGNVNHWLLNGGGWFFNLDYDKEILAKALAHKADELPLIIELVSKDKKYVICHADYP<br>FDEYEFGKPVDHQQVIWNRERISNSQNGIVKEIKGADTFIFGHTPAVKPLKFANQMYIDTGAVFCGNLTLIQV<br>QGEGA |
| **Gly-SspB-λ-phosphatase**<br>tetraglycine, $(GS)_4$, SspB (H. influenza, 5-114 Y11K A15E), $(GS)_4$, PPase ORF221 of bacteriophage lambda | GGGGGSGSGSGSSSPKRPKLLREYYDWLVDNSFTPYLVVDATYLGVNVPVEYVKDGQIVLNLSASATGNL<br>QLTNDFIQFNARFKGVSRELYIPMGAALAIYARENGDGVMFEPEEIYDELNIGSGSGSGSMRYYEKIDGSKY<br>RNIWVVGDLHGCYTNLMNKLDTIGFDNKKDLLISVGDLVDRGAENVECLELITFPWFRAVRGNHEQMMIDGL<br>SERGNVNHWLLNGGGWFFNLDYDKEILAKALAHKADELPLIIELVSKDKKYVICHADYPFDEYEFGKPVDHQ<br>QVIWNRERISNSQNGIVKEIKGADTFIFGHTPAVKPLKFANQMYIDTGAVFCGNLTLIQVQGEGA |
| **Gly-SspB-AuroraB**<br>tetraglycine, $(GS)_4$, SspB (H. influenza, 5-114 Y11K A15E), $(GS)_4$, human AuroraB (45-344) | GGGGGSGSGSGSSSPKRPKLLREYYDWLVDNSFTPYLVVDATYLGVNVPVEYVKDGQIVLNLSASATGNL<br>QLTNDFIQFNARFKGVSRELYIPMGAALAIYARENGDGVMFEPEEIYDELNIGSGSGSGSMSNVQPTAAPGQ<br>KVMENSSGTPDILTRHFTIDDFEIGRPLGKGKFGNVYLAREKKSHFIVALKVLFKSQIEKEGVEHQLRREIEIQ<br>AHLHHPNILRLYNYFYDRRRIYLILEYAPRGELYKELQKSCTFDEQRTATIMEELADALMYCHGKKVIHRDIKP<br>ENLLLGLKGELKIADFGWSVHAPSLRRKTMCGTLDYLPPEMIEGRMHNEKVDLWCIGVLCYELLVGNPPFE<br>SASHNETYRRIVKVDLKFPASVPMGAQDLISKLLRHNPSERLPLAQVSAHPWVRANSRRVLPPSALQSVA |
| human **INCENP** (834-902) | MDEAHPRKPIPTWARGTPLSQAIIHQYYHPPNLLELFGTILPLDLEDIFKKSKPRYHKRTSSAVWNSPPL |
